# PBS3 and EPS1 complete salicylic acid biosynthesis from isochorismate in Arabidopsis

**DOI:** 10.1101/601948

**Authors:** Michael P. Torrens-Spence, Anastassia Bobokalonova, Valentina Carballo, Christopher M. Glinkerman, Tomáš Pluskal, Amber Shen, Jing-Ke Weng

## Abstract

Salicylic acid (SA) is an important phytohormone mediating both local and systemic defense responses in plants. Despite over half a century of research, how plants biosynthesize SA remains unresolved. In Arabidopsis, a major part of SA is derived from isochorismate, a key intermediate produced by the isochorismate synthase (ICS), which is reminiscent of SA biosynthesis in bacteria. Whereas bacteria employ an isochorismate pyruvate lyase (IPL) that catalyzes the turnover of isochorismate to pyruvate and SA, plants do not contain an IPL ortholog and generate SA from isochorismate through an unknown mechanism. Combining genetic and biochemical approaches, we delineated the SA biosynthetic pathway downstream of isochorismate in Arabidopsis. We show that PBS3, a GH3 acyl adenylase-family enzyme important for SA accumulation, catalyzes ATP- and Mg^2+^-dependent conjugation of *L*-glutamate primarily to the 8-carboxyl of isochorismate and yields the key SA biosynthetic intermediate isochorismoyl-glutamate A. Moreover, EPS1, a BAHD acyltransferase-family protein with previously implicated role in SA accumulation upon pathogen attack, harbors a noncanonical active site and an unprecedented isochorismoyl-glutamate A pyruvoyl-glutamate lyase (IPGL) activity that produces SA from the isochorismoyl-glutamate A substrate. Together, PBS3 and EPS1 form a two-step metabolic pathway to produce SA from isochorismate in Arabidopsis, which is distinct from how SA is biosynthesized in bacteria. This study closes a major knowledge gap in plant SA metabolism and would help develop new strategies for engineering disease resistance in crop plants.

## Introduction

As sessile organisms, plants have evolved intricate chemical defense systems to perceive and defend against a plethora of pathogen invasions. Decades of research have established the role of SA as a pivotal phytohormone that signals plant defense responses in both local and distal tissues (Amick Dempsey, 1999). In addition to its function in defense signaling, SA also contributes to various aspects of plant growth and development, including photosynthesis, transpiration, ion uptake, thermogenesis, drought resistance and senescence (Raskin, 1992). Previous studies suggest that SA is biosynthesized from chorismate via two major metabolic routes (Dempsey et al., 2011). In Arabidopsis, a fraction of SA is produced from benzoic acid (BA) derived from general phenylpropanoid metabolism (León et al., 1995), whereas previous characterization of the Arabidopsis *sid2* mutant, defective in the *ICS1* gene (*AT1G74710*), revealed that the bulk of *de novo* biosynthesized SA upon *Pseudomonas syringae* infection is derived from isochorismate (Wildermuth et al., 2001). While SA-producing bacteria employ an IPL that directly converts isochorismate to SA, previous attempts to identify plant enzymes with *bona fide* IPL activity have not been successful (Zhou et al., 2018), suggesting an alternative mechanism for SA biosynthesis from isochorismate in plants (Chen et al., 2009).

Like *sid2*, additional Arabidopsis mutants, namely *pbs3* and *eps1*, exhibit impaired SA metabolism and susceptibility to bacterial pathogens (Nobuta et al., 2007; Jagadeeswaran et al., 2007; Lee et al., 2007; Zheng et al., 2009). *PBS3* (*GH3.12, AT5G13320*) encodes a member of the GH3 acyl-adenylate/thioester-forming enzyme family. Arabidopsis contains a total of 19 GH3 proteins with characterized members capable of catalyzing the conjugation *L*-amino acids to jasmonic acid, indole acetic acid and hydroxybenzoic acids, respectively (Westfall et al., 2012). Although its *in vivo* function remains unknown, PBS3 was shown to contain *in vitro* acyl acid amido synthetase activity, i.e., conjugating *L*-glutamate to 4-substituted BAs (Okrent et al., 2009; Westfall et al., 2012). Interestingly, SA appears to be a poor substrate of PBS3 and inhibits PBS3 activity *in vitro* (Okrent et al., 2009; Westfall et al., 2012). On the other hand, *EPS1* (*AT5G67160*) encodes a BAHD acyltransferase of unknown biochemical function. The *eps1* mutant phenotypically resembles *pbs3*, where both mutants display enhanced susceptibility to *P. syringae* and impaired *de novo* SA biosynthesis upon *P. syringae* infection (Zheng et al., 2009). Notably, EPS1 harbors an unusual active-site serine (Ser160) at the residue position corresponding to the highly conserved catalytic histidine in BAHD acyltransferases (Zheng et al., 2009; Levsh et al., 2016), suggesting it may contain unconventional catalytic activity different from the canonical acyltransferase activity (Supplemental Figure 1A, B and C). Together, these prior investigations suggest potential roles of PBS3 and EPS1 in SA metabolism (Chen et al., 2009).

In this study, we investigate the biochemical functions of PBS3 and EPS1 through untargeted metabolomics of various Arabidopsis SA-metabolism mutants (Zeilmaker et al., 2015; Zhang et al., 2017). Analysis of differentially accumulated metabolites between these mutant lines identified putative intermediates and products of SA metabolism that suggest possible catalytic roles of PBS3 and EPS1. *In vitro* enzyme assays and *de novo* reconstitution of SA biosynthesis in *Nicotiana benthamiana* established that PBS3 and EPS1 constitute the missing steps connecting isochorismate to SA production in Arabidopsis.

## Results

### *pbs3* and *eps1* are suppressors of the Arabidopsis *s3h*/*dmr6* double mutant

To resolve the SA biosynthetic pathway in plants, we first took a genetic approach. Two Arabidopsis 2-oxoglutarate/Fe (II)-dependent dioxygenases (AT4G10500 and AT5G24530) were recently characterized as the SA 3-hydroxylase (S3H) and SA 5-hydroxylase (S5H), respectively (Zhang et al., 2013; Zhang et al., 2017). The single null mutants of *S3H* and *S5H* in Arabidopsis, namely *s3h* and *dmr6*, exhibit various growth phenotypes and enhanced resistance to pathogens which are further exacerbated in the *s3h*/*dmr6* double mutant (van Damme et al., 2008; Zhang et al., 2013; Zeilmaker et al., 2015; Zhang et al., 2017) (Figure 1). Whereas the *s3h* plants accumulate SA at a similar level to wild-type plants, *dmr6* and particularly the *s3h/dmr6* double mutant contain significantly higher levels of free SA, SA 2-*O*-β-D-glucoside (SAG) and SA glucose ester (SGE) (Zhang et al., 2013; Zhang et al., 2017) (Figure 2A, Supplemental Figure 2, Supplemental Data 2). In addition to the over-accumulation of SA and SA-glucose conjugates, the *s3h*/*dmr6* double mutant displays a severe stunted-growth phenotype and significantly enhanced resistance against the virulent bacterial pathogen *P. syringae* pv. *tomato* DC3000 (*Pst*DC3000) (Zhang et al., 2017) (Figure 1). Collectively, S3H and S5H function as major SA catabolic enzymes that contribute to the proper maintenance of SA homeostasis in higher plants (Zhang et al., 2017). We therefore reasoned that *s3h*/*dmr6* provides an ideal genetic background to study Arabidopsis genes necessary for SA biosynthesis and signaling.

**Figure 1.**
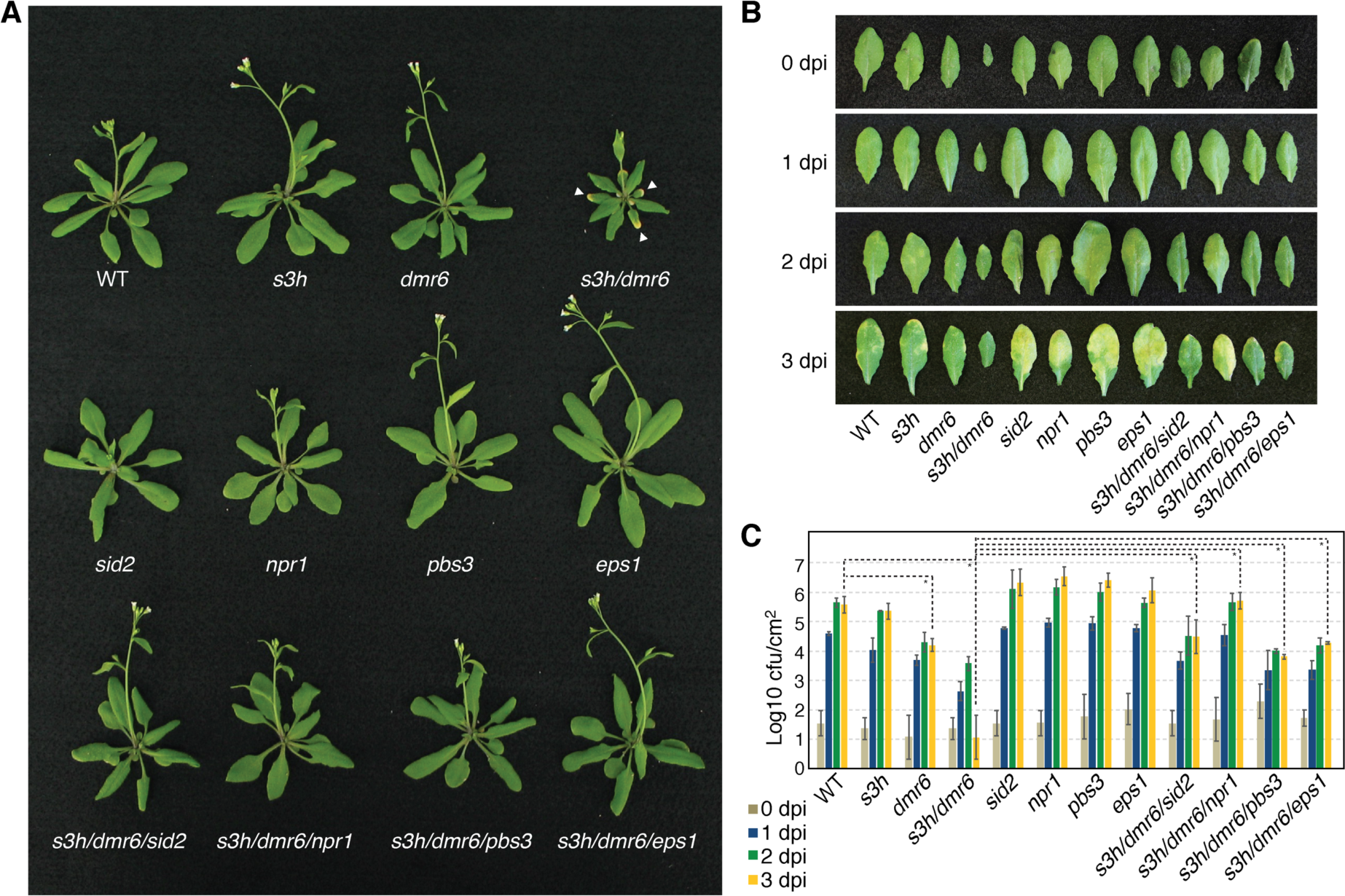
Characterization of the roles of *PBS3* and *EPS1* in plant growth and defense in an Arabidopsis mutant background that over-accumulates SA. **(A)** Four-week-old green-house-grown Arabidopsis plants. The *s3h/dmr6* double mutant exhibit stunted growth and early senescence (white arrows) phenotypes associated with hyper-accumulation of SA (Zhang et al., 2017), which could be rescued by crossing with the *sid2*, *npr1*, *pbs3* and *eps1* mutants, respectively. **(B)** Representative images of Arabidopsis rosette leaves infiltrated with a suspension of the *P. syringae* PstDC3000 strain. Dpi, days post-inoculation. **(C)** Quantification of the bacterial growth in Arabidopsis rosette leaves infiltrated with *P. syringae* PstDC3000 strain. Leaves were sampled at 0 through 4 dpi to measure the growth of the bacterial pathogen. The Logarithmic errors (δz) were calculated from the standard deviations of the raw data (n = 3) according to Baird (Baird, 1995). Statistical analysis was conducted by one-tailed unpaired *t*-test. \**P* < 0.05. Cfu, colony-forming unit.

**Figure 2.**
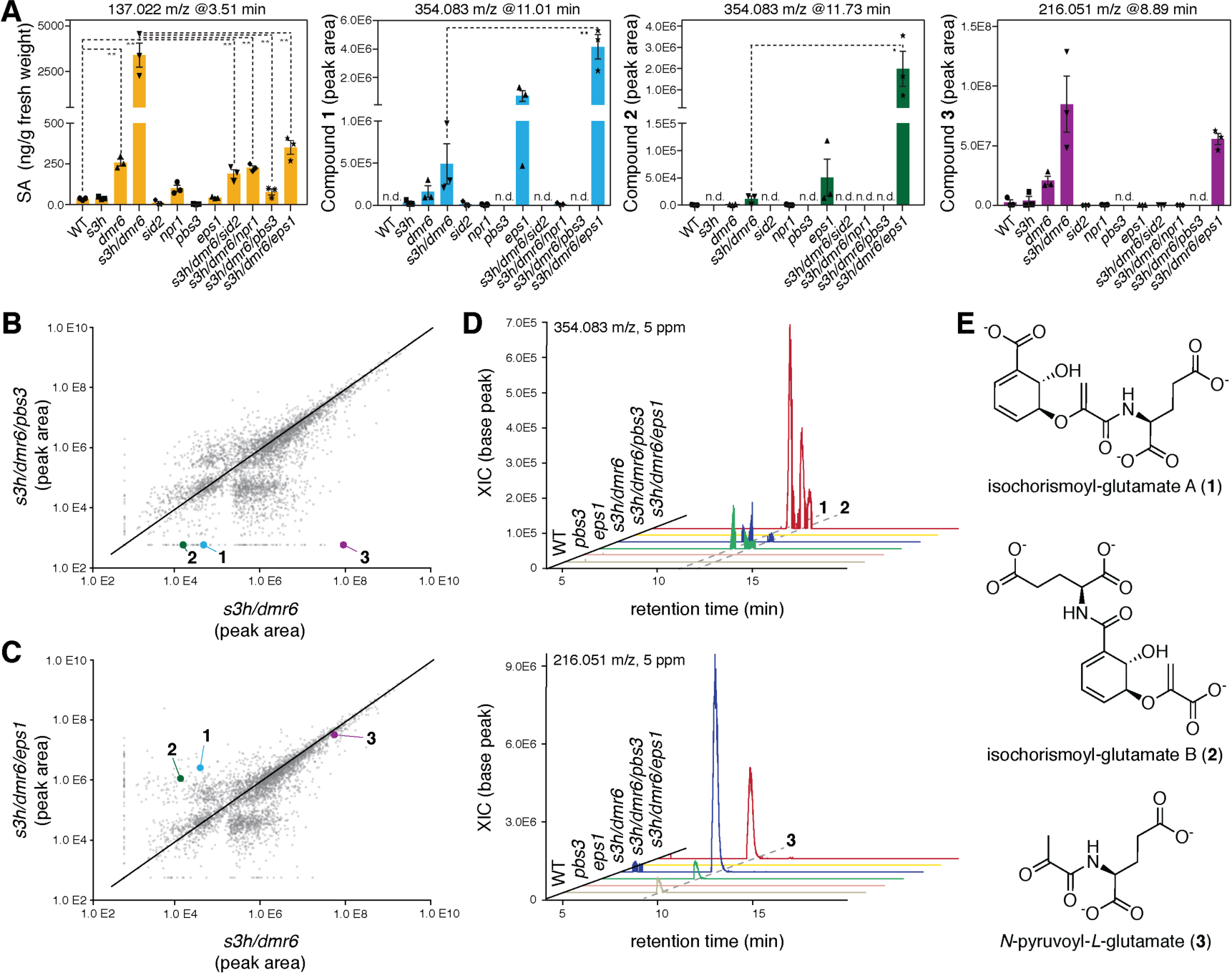
Key SA biosynthetic intermediates and products revealed by LC-HRAM-MS-based metabolic profiling of Arabidopsis mutants. **(A)** Quantification of SA and three newly identified metabolites (**1**-**3**) that are differentially accumulated across various Arabidopsis SA biosynthetic and signaling mutants (n = 3). Error bars indicate standard error of the mean (SEM) based on biological triplicates. Statistical analysis was conducted by one-tailed unpaired *t*-test. \*\**P* < 0.01, \**P* < 0.05. **(B)** Pairwise scatter plot comparing the abundance of global metabolic features of *s3h/dmr6* and *s3h/dmr6/pbs3*. The plot shows the accumulation of **1**-**3** in *s3h/dmr6*, which are absent in *s3h/dmr6/pbs3*. Data were collected using LC-HRAM-MS under negative mode. Each dot represents a metabolic feature defined by unique m/z and retention time values (n =3). **(C)** Pairwise scatter plot comparing the abundance of global metabolic features of *s3h/dmr6* and *s3h/dmr6/eps1*, which shows hyper-accumulation of **1** and **2** in *s3h/dmr6/eps1* compared to *s3h/dmr6* (n =3). **(D)** Representative extracted ion chromatograms (XICs) corresponding to **1**-**3** in select Arabidopsis mutants. The metabolites were resolved by hydrophilic interaction chromatography. **(E)** Structural assignment of **1**-**3** (also see Supplemental Figure 4A, B, Supplemental Figure 5).

To test this idea, we crossed *s3h*/*dmr6* with the extensively studied SA biosynthetic mutant *sid2* and SA signaling mutant *npr1 (Cao et al., 1997)*, and generated the corresponding *s3h*/*dmr6/sid2* and *s3h*/*dmr6/npr1* triple mutants, respectively. Both triple mutants display a complete rescue of the stunted-growth phenotype and a partial reversion of the pathogen resistance phenotype of the *s3h*/*dmr6* double mutant (Figure 1). Consistent with the recovered growth, both the *s3h*/*dmr6/sid2* and *s3h*/*dmr6/npr1* plants also exhibit significantly reduced levels of free SA, SAG and SGE compared to those of the *s3h*/*dmr6* plants (Figure 2A, Supplemental Figure 2). These observations indicate that hyper-accumulation of SA in the *s3h*/*dmr6* plants not only requires ICS activity, but also NPR1, one of the primary SA receptors in Arabidopsis (Wu et al., 2012; Ding et al., 2018). Upon the initial activation of SA-mediated defense response, NPR1 likely activates a positive feedback loop that further amplifies SA production by upregulating SA biosynthetic genes. These results also suggest that loss-of-function mutations in other genes involved in SA biosynthesis may also alleviate the mutant phenotype of *s3h*/*dmr6*.

To probe the functions of *PBS3* and *EPS1* in the *s3h*/*dmr6* genetic background, we crossed *pbs3* and *eps1* with *s3h*/*dmr6*, and generated the *s3h*/*dmr6/pbs3* and *s3h*/*dmr6/eps1* triple mutants, respectively. Like the *s3h*/*dmr6/sid2* and *s3h*/*dmr6/npr1* triple mutants, the *s3h*/*dmr6/pbs3* and *s3h*/*dmr6/eps1* triple mutants display a complete and nearly complete rescue of the stunted-growth phenotype, respectively (Figure 1A). The *s3h*/*dmr6/pbs3* and *s3h*/*dmr6/eps1* plants are also less resistant to *Pst*DC3000 compared to the *s3h*/*dmr6* plants (Figure 1B, C). The *s3h*/*dmr6/pbs3* and *s3h*/*dmr6/eps1* plants accumulate free SA, SAG and SGE at levels significantly less than those of the *s3h*/*dmr6* plants (Figure 2A, Supplemental Figure 2). These new genetic observations suggest that PBS3 and EPS1 may be directly involved in SA biosynthesis or participate in other auxiliary metabolic processes necessary for proper SA accumulation in Arabidopsis (Wildermuth et al., 2001).

### Untargeted metabolomics revealed key intermediates and side products of SA biosynthesis in Arabidopsis

To define the precise biochemical functions of PBS3 and EPS1 in plant, we performed untargeted metabolomic profiling of three-week-old rosette leaves from wild-type Arabidopsis and the aforementioned single and multiple mutants using liquid-chromatography high-resolution accurate-mass mass-spectrometry (LC-HRAM-MS) (Supplemental Data 1, 2). Comparative analysis of the resultant metabolomic datasets between specific mutant backgrounds identified three new metabolite features (**1**-**3**) that accumulate in the *s3h*/*dmr6* plants but are below the detection limit in the *pbs3* and *s3h*/*dmr6/pbs3* plants, relating them to PBS3 function (Figure 2, Supplemental Figure 3). Analysis of the mass spectra in the context of plausible isochorismate-derived SA biosynthetic pathways led to tentative structural assignment of these metabolites. **1** and **2** were resolved as two regioisomers of isochorismoyl-glutamate with glutamate conjugated to either the 8- or 1-carboxyl of isochorismate, referred to as isochorismoyl-glutamate A and B, respectively (Supplemental Figure 4). **3** was identified as *N*-pyruvoyl-*L*-glutamate, a possible degradation product of **1** (Supplemental Figure 5A). The identity of **3** was further confirmed by chemical synthesis (Supplemental Figure 5B, C, D). Since PBS3 reportedly contains *in vitro* acyl acid amido synthetase activity and prefers *L*-glutamate as its amino acid donor substrate (Okrent et al., 2009), our new findings suggest **1** and **2** may be the *in vivo* enzymatic products of PBS3, which are depleted in the *pbs3* mutant background. Moreover, **1** is likely further catabolized *in vivo* to yield **3**, and under such reaction scheme, the other co-product would be SA (Figure 3A). Intriguingly, the *eps1* and *s3h*/*dmr6/eps1* plants contain levels of **1** and **2** orders of magnitude higher than those of the wild-type and *s3h*/*dmr6* plants respectively (Figure 2A, C, D, Supplemental Figure 3B, D), suggesting that EPS1 may be involved in the catabolic pathway downstream of **1** and **2**.

**Figure 3.**
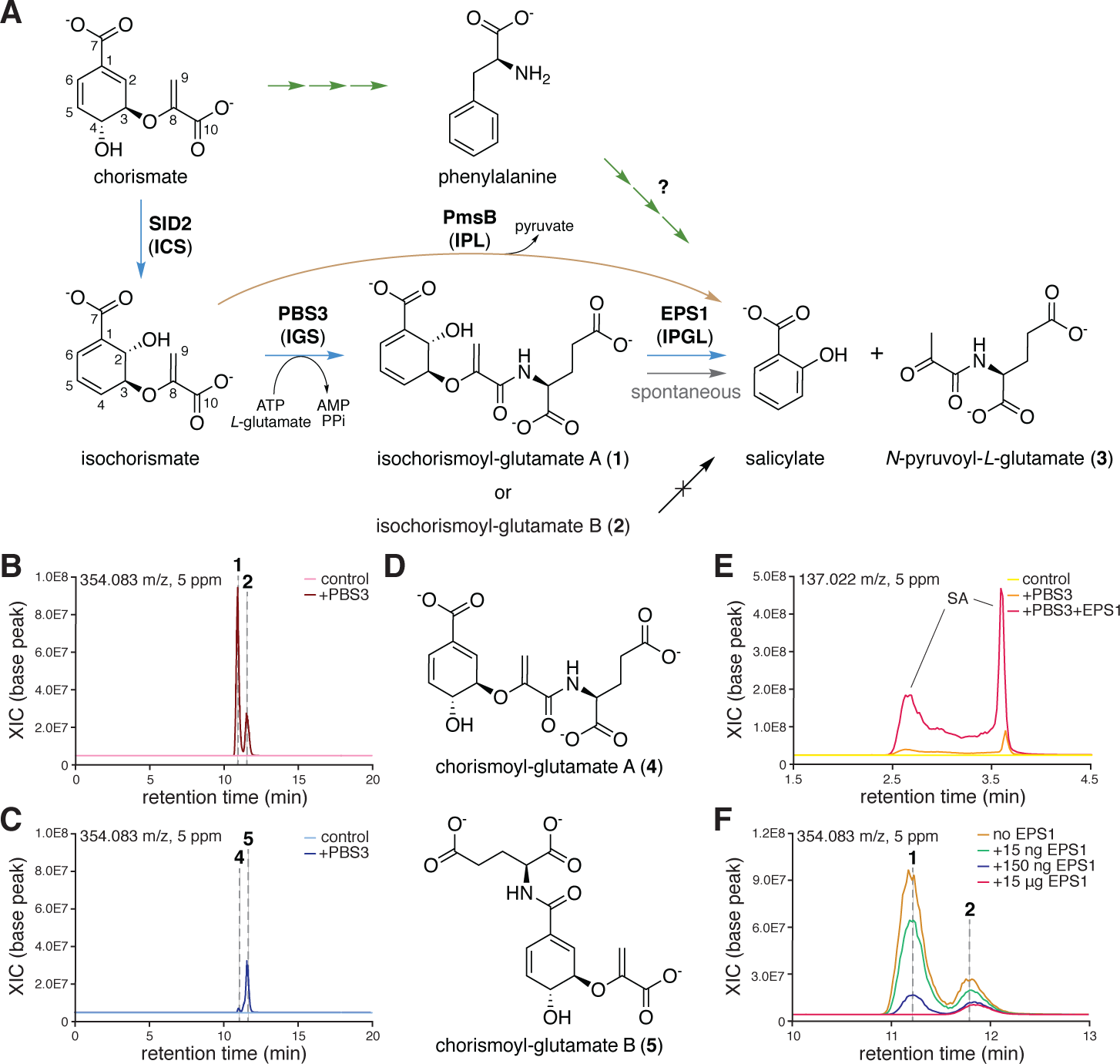
The biochemical functions of PBS3 and EPS1 in plant SA biosynthesis. **(A)** The SA biosynthetic pathways in higher plants. The isochorismate-derived pathway is highlighted by blue arrows. The phenylpropanoid-metabolism-derived pathway is highlighted by green arrows, which has not been fully resolved to date. Some SA-producing bacteria employ an IPL that directly converts isochorismate to SA, which is highlighted by a brown arrow. Enzyme names are in parentheses under the representative protein names. **(B)** Extracted ion chromatogram (XICs) showing the *in vitro* enzyme activity of PBS3 that produces **1** and **2** starting from a chorismate-SID2 preassay that contains approximately equal molar isochorismate and chorismate as potential acyl substrates. **(C)** PBS3 can also produce **4** and **5** from an assay buffer that contains only chorismate as the acyl substrate. The control traces in **b** and **c** represent enzyme assays performed without adding PBS3. **(D)** Structural assignment of **4** and **5** (also see Supplemental Figure 4C, D). **(E)** Representative XICs showing spontaneous decay of **1** that produces SA at the end of PBS3-isochorismate enzyme assay (orange trace). The addition of EPS1 (15 µg) converts all remaining **1** to SA within 5 min (red trace). The chromatograms shown were resolved using hydrophilic interaction chromatography. SA splits into two peaks under such chromatographic condition. **(F)** Representative XICs showing EPS1 preferably turns over **1** to produce SA and **3** in a EPS1 enzyme-concentration-dependent manner (also see Supplemental Figure 9C, D). EPS1 also exhibits minor activity that turns over **2**.

### The *in vitro* enzymatic functions of PBS3 and EPS1

We next characterized the *in vitro* biochemical functions of PBS3 and EPS1 using purified recombinant proteins in enzyme assays. Since isochorismate is not commercially available, we devised a pre-assay to enzymatically synthesize isochorismate from chorismate using recombinant SID2. Starting with 5 mM chorismate, the reaction reaches equilibrium with approximately equal molar ratio of isochorismate and chorismate at the end of the pre-assay (Supplemental Figure 6A). Consistent with the findings from the Arabidopsis metabolomics experiments, when PBS3 was added to the pre-assay containing approximately equal molar amounts of isochorismate and chorismate, it exhibits specific ATP- and Mg^2+^-dependent acyl acid amido synthetase activity (Supplemental Figure 6B) towards isochorismate with *L*-glutamate as the preferred amino acid substrate (Supplemental Figure 6C, D), and yields **1** as the main enzymatic product together with **2** as a minor product (Figure 3B and Supplemental Figure 7). In the absence of isochorismate, PBS3 also exhibits acyl acid amido synthetase activity towards chorismate and produces regio-isomeric chorismoyl-glutamate A and B (**4** and **5** respectively, Figure 3C, D, Supplemental Figure 4), although these products were not detected in Arabidopsis metabolomics datasets. Therefore, PBS3 functions as a substrate- and regio-selective isochorismoyl-glutamate synthase (IGS) both *in vivo* and *in vitro*.

A previous study of the crystal structure of PBS3 in complex with SA and adenosine monophosphate (AMP) revealed that SA binds to PBS3 in a catalytically nonproductive pose with its carboxyl group pointing away from the catalytic center (Westfall et al., 2012) (Supplemental Figure 8A). We performed docking simulation of isochorismate and chorismate to the PBS3 structure (PDB: 4EQL) (Westfall et al., 2012). The 1-carboxyl of isochorismate preferably adopts a similar orientation as the SA carboxyl, which in turn presents the isochorismate 8-carboxyl in a catalytically productive conformation towards the catalytic center, readying it for adenylation (Supplemental Figure 8B). Although chorismate could be docked to PBS3 active site in a similar orientation as isochorismate, it appears to be a less favored substrate due to the differential placement of its 4-hydroxyl (Supplemental Figure 8C). These ligand docking results reveal features of the PBS3 active site consistent with the experimentally observed substrate- and regio-selectivity of PBS3.

We noticed that **1** produced by PBS3 in enzyme assays is unstable and spontaneously decays to produce SA and **3** at a rate of 0.21 pkat (Figure 3E, Supplemental Figure 9A, B), whereas **2** is stable under our assay conditions. Interestingly, addition of EPS1 to the PBS3 enzyme assay resulted in rapid turnover of **1** to SA and **3**, with the catalyzed rate four orders of magnitude faster than the spontaneous decay rate of **1** under our assay condition (Figure 3E, Supplemental Figure 5C, Supplemental Figure 9C, D). EPS1 also displays minor activity to turn over **2** (Figure 3F), but no measurable lyase or hydrolase activity against isochorismate, chorismate, **3**, **4** and **5** was observed, suggesting its IPGL activity is selective towards **1**. Altogether, our *in vitro* enzyme assays further confirmed the chemical identities of **1**-**3**, established the specific acyl acid amido synthetase activity of PBS3 in the production of the key SA biosynthetic intermediate **1**, and revealed that EPS1 is an unprecedented IPGL that significantly enhances the rate of SA production from **1**.

### Reconstitution of *de novo* SA biosynthesis in *N. benthamiana*

Upon pathogen attack, Arabidopsis rapidly activates *de novo* SA biosynthesis in both local and systemic tissues, mostly through the isochorismate-derived pathway (Wildermuth et al., 2001). To identify the core Arabidopsis gene set necessary and sufficient for this rapid *de novo* SA biosynthesis, we sought to reconstitute *de novo* SA biosynthesis in *N. benthamiana* using the Agrobacterium-mediated transient transgenic expression technique (Sainsbury et al., 2009) (Figure 4A). Infiltration of *N. benthamiana* leaves with Agrobacterium carrying empty vector did not trigger significant level of *de novo* SA biosynthesis compared to the control leaves, thus providing a clean background for this reconstitution exercise (Supplemental Figure 10A). Experimenting with several genes previously implicated in SA metabolism, we discovered that coexpression of *SID2* together with *SID1* (*AT4G39030*), a chloroplast-envelope-localized MATE-family transporter required for pathogen-induced SA accumulation in Arabidopsis (Nawrath et al., 2002), resulted in ectopic accumulation of isochorismate in *N. benthamiana* leaves (Figure 4B, Supplemental Figure 10B, Supplemental Figure 11). Furthermore, coexpression of *PmsB*, a bacterial *IPL* (Verberne et al., 2000), together with *SID1* and *SID2* is sufficient to elicit *de novo* SA biosynthesis in *N. benthamiana* leaves (Figure 3A, Figure 4B, Supplemental Figure 12). In comparison, coexpression of *PBS3* together with *SID1* and *SID2* resulted in not only *de novo* production of SA but also a depletion of isochorismate level in *N. benthamiana* leaves (Figure 4B, Supplemental Figure 12). The *PBS3-SID1-SID2* coexpression also led to ectopic accumulation of **1**-**3**, which were absent in the *PmsB-SID1-SID2* coexpression experiment (Figure 4C, D, Supplemental Figure 12). In contrast, coexpression of *EPS1* together with *SID1* and *SID2* did not result in increased *de novo* production of SA, **1**, **2** and **3** nor any decrease of isochorismate level (Figure 4B, D). When *EPS1* was coexpressed together with *PBS3*, *SID1* and *SID2*, the levels of isochorismate, **1** and **2** were significantly reduced compared to those of the *PBS3-SID1-SID2* coexpression, although the accumulating levels of SA and **3** did not increase further (Figure 4B, D). Expression of *SID1*, *SID2*, *PmsB*, *PBS3* or *EPS1* alone did not elicit *de novo* SA synthesis nor yield measurable levels of any SA biosynthetic intermediates in *N. benthamiana* leaves (Figure 4B, D, Supplemental Figure 10B, Supplemental Figure 12). These reconstitution experiments in *N. benthamiana* further reinforce the critical function of *PBS3* in the plant isochorismate-derived SA biosynthetic pathway, and suggest *PBS3*, *SID1* and *SID2* as a minimum gene set necessary and sufficient to reconstitute *de novo* SA biosynthesis in a heterologous plant host. Although the *PBS3-EPS1-SID1-SID2* coexpression reduced the accumulation of **1** and **2** compared to *PBS3-SID1-SID2,* the addition of *EPS1* did not significantly enhance SA and **3** production further in *N. benthamiana* (Figure 4C, E). This could be attributed to the spontaneous decay of **1** over the course of the Agrobacterium infiltration experiment (5 days), existing IPGL activity in *N. benthamiana*, feedback inhibition of PBS3 by the *de novo* biosynthesized SA, and/or *N. benthamiana* endogenous SA production triggered by the initial SA accumulation resulted from the expression of Arabidopsis transgenes.

**Figure 4.**
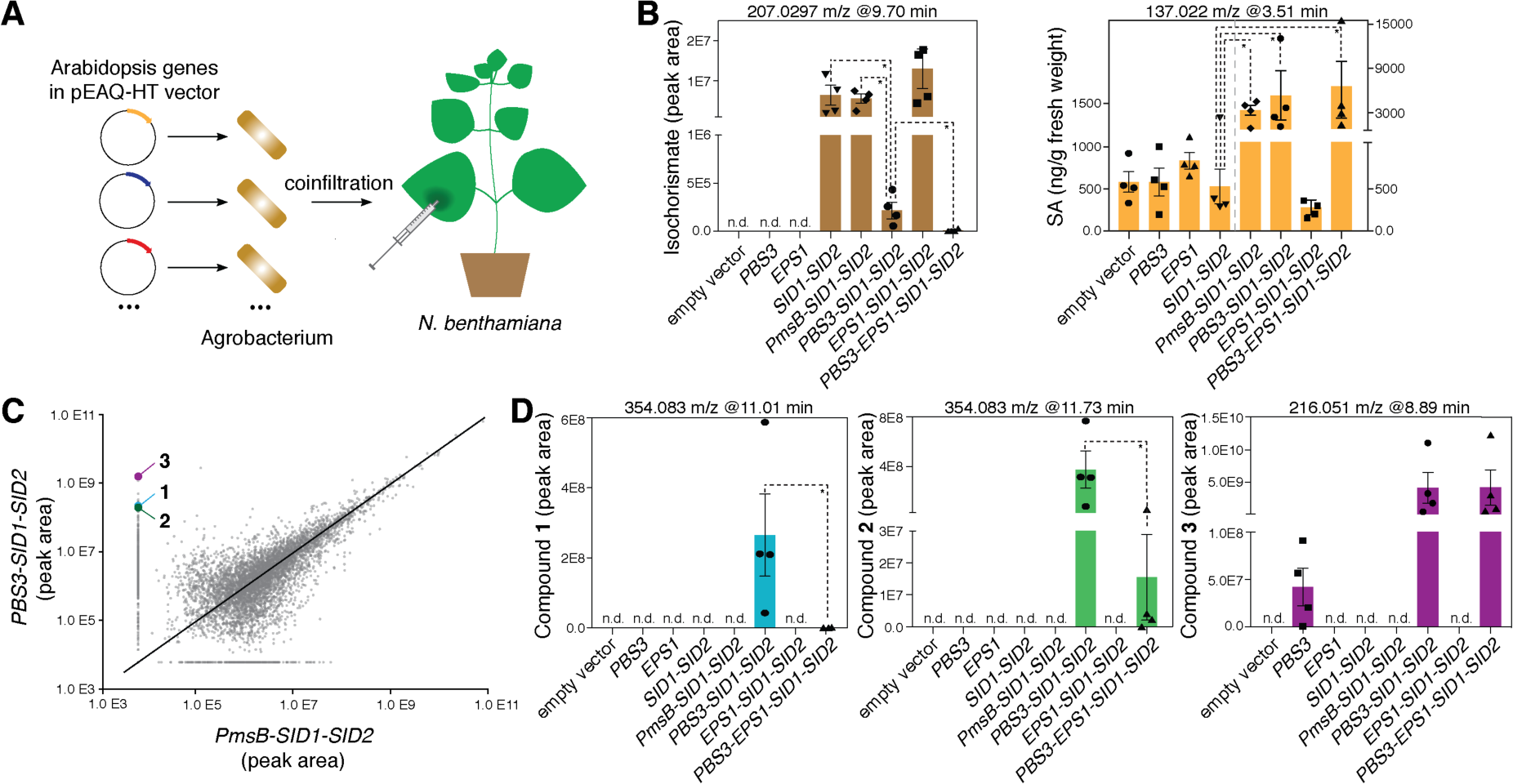
Reconstitution of *de novo* SA biosynthesis in *N. benthamiana* by Agrobacterium-mediated transient coexpression of transgenes. **(A)** Individual Arabidopsis genes were cloned into the pEAQ-HT vector (Sainsbury et al., 2009) and separately transformed into Agrobacterium. Combination of the resultant Agrobacterium strains was used to coinfiltrate the leaves of *N. benthamiana* plants. LC-HRAM-MS-based metabolic profiling of leaf extracts was performed 5 days post infiltration. **(B)** Relative abundance of isochorismate and SA across various coinfiltration experiments (n = 4). **(C)** Pairwise scatter plot comparing the abundance of global metabolic features of *PBS3-SID1-SID2* coexpression and *PmsB-SID1-SID2* coexpression in *N. benthamiana* leaves. The plot shows hyper-accumulation of **1**-**3** in *PBS3-SID1-SID2*-coinfiltrated leaves which are absent in the *PmsB-SID1-SID2* coexpression experiment. Data were collected using LC-HRAM-MS under negative mode. Each dot represents a metabolic feature defined by unique m/z and retention time values (n =4). **(D)** Relative abundance of **1**-**3** across various coinfiltration experiments (n = 4). Error bars indicate SEM based on biological quadruplicates. n.d., not detected. Statistical analysis was conducted by one-tailed Mann-Whitney test. \**P* < 0.05.

## Discussion

SA plays a central role in defense signaling of higher plants (Chen et al., 2009). In this study, we elucidated the previously elusive isochorismate-derived SA biosynthetic pathway in Arabidopsis (Figure 3A). PBS3 is an essential enzyme in this pathway immediately downstream of ICS, responsible for the production of key SA intermediate isochorismoyl-glutamate A. The important role of PBS3 in SA biosynthesis in higher plants is corroborated by a recent phylogenomic study showing that *PBS3* orthologs are present in several model flowering plants with predicted emergence predating the monocot/eudicot split (Okrent and Wildermuth, 2011). Isochorismoyl-glutamate A may be a conserved SA biosynthetic intermediate in flowering plants. Indeed, our findings are consistent with a recent independent study by Rekhter et al. describing the function of PBS3 in SA biosynthesis (Rekhter et al., 2019). The role of PBS3 in plant SA biosynthesis illustrates another case of the involvement of GH3 acyl adenylase-family enzyme in phytohormone metabolism and signaling. In other described cases, JAR1 (GH3.11) plays a role in the biosynthesis of the active form of jasmonate hormone JA-Ile (Westfall et al., 2012), whereas VAS2 (GH3.17) and WES1 (GH3.5) are responsible for attenuating the hormonal activity of auxin indole-3-acetic acid (IAA) and SA by conjugating them to amino acids (Park et al., 2007; Zheng et al., 2016; Westfall et al., 2016). Interestingly, PBS3 is involved in SA biosynthesis, meanwhile is also inhibited by SA, providing a feedback mechanism to regulate the metabolic flux into the isochorismate-branch of the SA biosynthetic pathway in plants (Okrent et al., 2009). Previous studies also suggest a pathogen-induced *SID2*-dependent and *PBS3*-independent pathway for SA biosynthesis may exist in Arabidopsis (Nobuta et al., 2007). It is likely that other members of the GH3 family may contain dedicated or promiscuous IGS activity that function in SA biosynthesis in addition to PBS3.

In contrast to PBS3, EPS1 belongs to a unique clade of BAHD acyltransferases only associated with the Brassicaceae family plants (Supplemental Figure 1C, Supplemental Data 5, Supplemental Data 6), indicating that EPS1 was recently recruited to the SA biosynthetic pathway in Brassicaceae. This evolutionary development could be driven by strong selection pressure to improve SA production kinetics amid pathogen attack in common ancestors of the extant Brassicaceae plants. EPS1 harbors an unusual active site reconfigured from an ancestral BAHD acyltransferase to catalyze the unprecedented IPGL chemistry (Supplemental Figure 1B), and thus presents an attractive case for future studies of molecular evolution that gave rise to novel catalytic function based on ancestral enzyme fold (Weng and Noel, 2012). Although loss of *EPS1* in Arabidopsis led to ectopic hyper-accumulation of isochorismoyl-glutamate A and B and compromised SA production, the *s3h*/*dmr6/eps1* triple mutant accumulates *N*-pyruvoyl-*L*-glutamate at a level comparable to that of the *s3h*/*dmr6* double mutant and an SA level higher than that of the *s3h*/*dmr6/pbs3* triple mutant (Figure 2A), suggesting spontaneous decay of isochorismoyl-glutamate A does occur *in vivo* at the absence of EPS1. The chemical mechanism of isochorismoyl-glutamate A spontaneous decay into SA and *N*-pyruvoyl-*L*-glutamate was recently proposed (Rekhter et al., 2019). Furthermore, decay of isochorismate to SA or another PBS3-independent biosynthetic route could also contribute to the background SA accumulation in the *sid2* or *pbs3* mutants. It is also worth noting that *EPS1* does not appear to coexpress with other known SA-pathway genes in Arabidopsis, which further suggests its auxiliary role in SA biosynthesis. Non-Brassicaceae plants may either rely on the spontaneous decay of isochorismoyl-glutamate A alone or have independently evolved alternative IPGLs to assist SA production.

Our study fills the long-sought-after missing steps of the isochorismate-dependent SA biosynthetic pathway in Arabidopsis; however, several important aspects of SA metabolism and regulation remain to be addressed in plants. For instance, the enzymatic steps downstream of phenylalanine ammonia lyase (PAL) in the phenylpropanoid-derived SA biosynthetic branch remain unresolved (Figure 3A). Although a crippled isochorismate-derived SA biosynthetic branch led to significantly reduced SA accumulation in Arabidopsis, previous studies showed that suppression of PAL expression in tobacco (Pallas et al., 1996) and Arabidopsis (Huang et al., 2010) also causes major reduction in SA accumulation, suggesting that the regulatory mechanism and functional significance of the partition of these two major SA biosynthetic pathway branches under different physiological and pathogen-attack conditions in different plant hosts are yet to be elucidated. A full understanding of SA metabolism in the plant kingdom will help device comprehensive strategies to engineer disease resistance in crops.

## Methods

### Plant materials and growth conditions

*Arabidopsis thaliana* and *Nicotiana benthamiana* plants were grown at 22 °C in a greenhouse under long-day condition (16-h light/8-h). The Arabidopsis T-DNA insertional null mutants *s3h*, *dmr6-2*, *sid2-4*, *npr1-1*, *pbs3-2*, and *eps1-2* in Columbia background have been previously characterized to be null mutants (Cao et al., 1997; Zheng et al., 2009; Zhang et al., 2013; Zeilmaker et al., 2015), and correspond to the Arabidopsis Biological Resource Center (ABRC) lines under the accessions SALK_059907, GK_249H03, SALK_133146, CS3726, SALK_018225C, and SAIL_734_F07, respectively (Alonso, 2003; Kleinboelting et al., 2012). For simplicity, the mutants were mentioned throughout the paper without their specific allele names called out.

### Pseudomonas syringae infection assay

*P. syringae* inoculations were performed according to previously described protocol (Katagiri et al., 2002). Briefly, *Pst* DC3000 was streaked out from a –80 °C glycerol stock onto King’s medium B (Protease Peptone #3 20 g/L, glycerol 10 mL/L, K_2_HPO_4_ 1.5 g/L, MgSO_4_·7H_2_O 1.5 g/L, Bacto agar 15 g/L, ph 7.2) with 50 mg/mL rifampicin, and grown for 2 days at 30 °C. Fresh bacteria was transferred to a 3 mL liquid culture and grown overnight at 30 °C, a final 200 mL liquid culture was grown overnight until final OD_600_ = 0.8. The culture was pelleted, and diluted in sterile water to a final concentration of 1 × 10^6^ colony-forming unit (cfu)/mL. 6-week-old Arabidopsis leaves were infiltrated with the bacterial suspension on their abaxial side with a 1 mL needleless syringe. At 0, 1, 2, 3 dpi, Infected leaves were harvested and surface sterilized for 1 minute with 70% ethanol and then rinsed with distilled water. Two leaf disks for each genotype and each triplicate were cut with a 0.3 cm^2^ cork borer. Leaf disks were placed in a 2.0 mL microfuge tube with 100 µL sterile distilled water, and thoroughly macerated using a plastic pestle. Ground tissue was diluted up to 1 mL with sterile water, and mixed by vortexing. Four serial dilutions were performed and blotted onto King’s medium B in 10 µL triplicate drops. After incubation at 30°C for two days, colonies were counted to derive cfu values.

### Transient expression of genes in *N. benthamiana*

The open reading frame (ORF) of target genes to be expressed were cloned into pEAQ-HT (Sainsbury et al., 2009). Vectors were transformed into *Agrobacterium tumefaciens* strain LBA4404 (Hoekema et al., 1983). Bacteria were cultivated at 30 °C in YM Medium (0.4 g/L yeast extract, 10 g/L mannitol, 0.1 g/L NaCl, 0.2 g/L MgSO_4_·7H_2_O, 0.5 g/L K_2_HPO_4_·3H_2_O) to an OD_600_ between 2.0 and 2.5, washed with 0.5X PBS, then diluted with 0.5X PBS to an OD_600_ of 0.3. For co-infiltration of multiple gene constructs, *A. tumefaciens* strains were mixed in equal concentrations to a final OD_600_ of 0.3. 0.5 mL of bacterial suspension was infiltrated into leaves of *N. benthamiana* plants between 4 and 6 weeks of age. Leaves were harvested 5 days post infiltration for untargeted metabolomics analysis.

### Metabolomic profiling of plant extracts

For untargeted metabolomics analysis, 3-week-old Arabidopsis rosette leaves or *N. benthamiana* leaves 5 days post *A. tumefaciens* infiltration were harvested, suspended in 50% methanol (500 µL per 100 mg fresh weight) and disrupted using a TissueLyser (Qiagen) and zirconia beads. Tissue debris were removed by centrifugation. 2 µL injections of the clarified extracts were separated using a 150 x 2.1 mm ZIC-pHILIC column (5 µm particle size, EMD Millipore). The chromatographic gradient was run at a flow rate of 0.15 mL/min as follows: 0–20 min, linear gradient from 80% to 20% solvent B; 20–20.5 min, linear gradient from 20% to 80% solvent B; 20.5–28 min, hold at 80% solvent B. Solvent A was 20 mM ammonium carbonate, 0.1% ammonium hydroxide; solvent B was acetonitrile. The column oven and autosampler tray were held at 25 °C and 4 °C, respectively. Data was collected with a Q-Exactive Orbitrap mass spectrometer (Thermo Scientific) for MS^1^ scans and dd-MS^2^ (data-dependent MS/MS) in both positive and negative polarity modes. The various arabidopsis leaf extracts were also additionally run through a C18 column (Kinetex 2.6-µm, 100-Å, 150 × 3 mm) with a gradient of Solvent A (0.1% formic acid in H_2_O) and Solvent B (0.1% formic acid in acetonitrile; 5% B for 2 min, 5-80% B over 40 min, 95% B for 4 min, and 5% B for 5 min; flow rate 0.8 mL/min) before detection by the Q-Exactive mass spectrometer. Full MS resolution, 70,000; mass range, 100– 1,250 m/z; dd-MS2 resolution, 17,500; loop count, 5; collision energy, 15–35 eV (stepped); dynamic exclusion, 1 s. The raw data were processed using MZmine 2 (Pluskal et al., 2010) and further analyzed using metaboanalyst (Chong et al., 2018). Statistical analysis was conducted using Prism 8.

### Chemical synthesis

#### General methods

All reactions were performed under nitrogen unless otherwise noted. All reagents and solvents were used as supplied without further purification unless otherwise noted. Column chromatography was conducted using Silicycle SiliaFlash P60 SiO_2_ (40–63 µm). Analytical TLC was conducting using Millipore SiO_2_ 60 F_254_ TLC (0.250 mm) plates. HPLC was conducted using a pair of Shimadzu LC-20AP pumps, a Shimadzu CBM-20A communications module, a Shimadzu SPD-20A UV-Vis detector, and a Phenomenex Kinetex 5u C18 100 Å Axia column (150 x 21.2 mm). Melting points were obtained using a Mel-Temp II apparatus and are uncorrected. ^1^H and ^13^C NMR spectra were obtained using a Bruker Avance Neo 400 MHz spectrometer equipped with a 5 mm Bruker SmartProbe. IR spectra were obtained using a Bruker Alpha 2 with a Platinum ATR accessory. Mass spectrometric analysis was performed on a JEOL AccuTOF-DART.

#### Di-*tert*-butyl *N*-pyruvoyl-*L*-glutamate (**S2**) (Koji et al., 2008)

A vigorously stirred solution of pyruvic acid (0.35 mL, 5.1 mmol) in CH_2_Cl_2_ (35 mL) at 23 °C was treated sequentially with EDCI (790 mg, 5.1 mmol) and HOBt (690 mg, 5.1 mmol). The reaction mixture was allowed to stir at 23 °C for 5 min, at which time protected glutamate **S1** (1.0 g, 3.4 mmol) was added and the reaction was stirred at 23 °C for an additional 24 h. After 24 h, the reaction mixture was triturated with Et_2_O (300 mL) and filtered through Celite. The filtrate was concentrated on a rotary evaporator and the residue was purified by flash chromatography (SiO_2_, 20% EtOAc/hexanes) to provide **S2** as a white solid (240 mg, 22%): mp 64–66 °C; ^1^H NMR (CDCl_3_, 400 MHz) δ 7.44 (d, *J* = 8.4 Hz, 1H), 4.40 (td, *J* = 8.2, 4.8 Hz, 1H), 2.43 (s, 3H), 2.32–2.19 (m, 2H), 2.18–2.10 (m, 1H), 2.00–1.89 (m, 1H), 1.44 (s, 9H), 1.40 (s, 9H); ^13^C NMR (CDCl_3_, 100 MHz) δ 196.2, 171.8, 170.1, 160.0, 82.8, 80.9, 52.3, 31.4, 28.1, 28.0, 27.5, 24.5; IR (film) ν_max_ 3365, 1741, 1714, 1671, 1512, 1367, 1258, 1230, 1144, 844, 620, 606 cm^−1^; HRMS (DART-TOF) *m/z* 330.1902 (C_16_H_27_NO_6_ + H^+^ requires 330.1917).

#### *N*-pyruvoyl-*L*-glutamate (**3**) (Wehbe et al., 2004)

A vigorously stirred suspension of **S2** (50 mg, 0.15 mmol) in THF (1.2 mL) at 23 °C was treated with 3 N HCl(aq) (3.0 mL). The reaction mixture was allowed to stir at 23 °C for 24 h, then concentrated under a stream of N_2_. The resulting solid residue was purified by preparative reverse-phase HPLC (CH_3_CN-0.1% TFA/H_2_O-0.1% TFA 5:95 to 95:5 over 30 min, 7 mL/min, *R*_t_ = 10.1 min) to provide **3** as a hygroscopic colorless film (16 mg, 48%): ^1^H NMR (DMSO-*d*_6_, 400 MHz) δ 12.47 (bs, 2H), 8.68 (d, *J* = 8.1 Hz, 1H), 4.23 (ddd, *J* = 9.6, 8.1, 4.7 Hz, 1H), 2.34 (s, 3H), 2.26 (t, *J* = 7.4 Hz, 2H), 2.06 (dtd, *J* = 15.5, 7.1, 4.7 Hz, 1H), 1.91 (ddt, *J* = 14.0, 9.7, 7.1 Hz, 1H); ^13^C NMR (DMSO-*d*_6_, 100 MHz) δ 196.7, 173.9, 172.4, 161.2, 51.4, 30.1, 25.6, 24.9; IR (film) ν_max_ 2926, 1719, 1637, 1524, 1411, 1358, 1165, 969, 611 cm^−1^; HRMS (DART-TOF) *m/z* 218.0651 (C_8_H_11_NO_6_ + H^+^ requires 218.0665).

### Recombinant protein production and purification

*E. coli* cultures transformed with coding sequence containing pHis-8-4 expression vectors were grown at 37 °C in TB media to an optical density of 1.5, induced with 1 mM isopropyl-*β*-*D*-thiogalactoside (IPTG), and allowed to grow for an additional 16 h at 18 °C. *E. coli* cells were harvested by centrifugation, washed with phosphate buffered saline, resuspended in 100 mL of lysis buffer [(50 mM phosphate buffer pH 6.2, 0.5 M NaCl, 20 mM imidazole, and 2 mM dithiothreitol (DTT)], and lysed with five passes through a M-110L microfluidizer (Microfluidics). The resulting crude protein lysate was clarified by centrifugation prior to Qiagen Ni-NTA gravity flow chromatographic purification. After loading the clarified lysate, the protein bound Ni-NTA resin was washed with 10 column volumes of lysis buffer and eluted with 1 column volume of elution buffer (50 mM phosphate buffer pH 6.2, 0.5 M NaCl, 200 mM imidazole, and 2 mM DTT). Purified recombinant enzymes were concentrated and washed multiple times with storage buffer (25 mM phosphate buffer pH 6.2, 50 mM NaCl, and 2 mM DTT) using an amicon ultra-15 centrifugal filter unit with ultracel-30 membrane (EMD Millipore) to achieve a final protein concentration of 10 mg/mL.

### Enzyme assays

Initial isochorismate synthase activity titration assays were performed in 100 µL reactions composed of 25 mM Tris and 5 mM chorismate. Reactions were started with the addition of 0.5 µg recombinant SID2, incubated at 25 °C for various time points, and quenched with the addition of 100 µL MeOH. To establish PBS3 activity *in vitro*, isochorismate was first generated enzymatically by adding 2 µg of SID2 to a 100 µL reaction containing 50 mM Tris, 1 mM *L*-glutamate, 5mM chorismate, 5 mM ATP, 5 mM MgCl_2_ and incubating for 5 minutes at 25 degrees. After incubation, 0.8 µg of PBS3 was added to each reaction mixture, incubated at 25 °C for various time points and then quenched with 100 µL MeOH. The ATP- and Mg^2+^ dependence of PBS3 was tested with identical reactions excluding ATP and MgCl_2_, respectively. 1 mM *L*-alanine, *L*-arginine, *L*-asparagine, *L*-aspartic acid, *L*-cysteine, *L*-glutamine, glycine, *L*-histidine, *L*-isoleucine, *L*-leucine, *L*-lysine, *L*-phenylalanine, *L*-proline, *L*-serine, *L*-threonine, *L*-tryptophan, *L*-tyrosine or *L*-valine was substituted for the *L*-glutamate in separate assays to assess the proteinogenic *L*-amino acid substrate selectivity of recombinant PBS3. The relative amino acid preference was determined by measuring AMP production. To characterize the biochemical function of EPS1, isochorismoyl-glutamate conjugates were first produced enzymatically in 100 µL reaction systems containing 50 mM Tris, 1 mM *L*-glutamate, 5mM chorismate, 5 mM ATP, and 5 mM MgCl_2_ with addition of 2 µg of SID2 and 8 µg of PBS3 incubated for 20 minutes at 25 °C. Various amounts of recombinant EPS1 and were added to the reactions, incubated for an additional 5 minutes at 25 °C prior to quenching with an equal volume of MeOH. Enzyme assays were analyzed by LC-HRAM-MS using ZIC-pHILIC column following the same procedure as described above.

### Sequence alignment, phylogenetic analysis and homology modeling

Sequence alignment shown in Supplemental Figure 1A was generated using T-coffee (Di Tommaso et al., 2011), and displayed using ESPript 3.0 (Robert and Gouet, 2014). The Neighbor-Joining (NJ) tree shown in Supplemental Figure 1A and Supplemental Data 4 was inferred using MEGA 7 (Kumar et al., 2016), based on sequence alignment built using MUSCLE (Edgar, 2004) (Supplemental Data 5). The bootstrap percentages were based on 1000 replications. The EPS1 structural homology model was generated using the SWISS-MODEL (Waterhouse et al., 2018) server with Arabidopsis HCT (PDB: 5KJU) as the template.

### Ligand docking

Docking simulation of chorismate and isochorismate into the active site of Arabidopsis PBS3 crystal structure (PDB: 4EQL) (Westfall et al., 2012) was performed using AutoDock Vina 1.1.2 (Trott and Olson, 2010). The bound ligands in 4EQL were removed before docking. The docking simulation generated top nine possible poses for each of the docked ligands, which were ranked by the predicted affinity (kcal/mol). The top poses for isochorismate and chorismate were shown in Supplemental Figure 8B and c respectively. Molecular graphics were rendered with PyMOL 2.0.7 (Schrödinger).

## Acknowledgement

We thank the Bioinformatics and Research Computing and the Metabolite Profiling core Facilities at the Whitehead Institute for assistance with the metabolomics experiments and data analysis. We thank Zhixiang Chen for providing numerous Arabidopsis SA-pathway mutants. We thank Jen Sheen for providing the *P. syringae* pv. *tomato* DC3000 strain. This work was supported by the Pew Scholar Program in the Biomedical Sciences, the Searle Scholars Program and the National Science Foundation (CHE-1709616).

## Author Contributions

M.P.T.-S. and J.-K.W. designed the research. M.P.T.-S. performed the cloning, protein expression, biochemical assays, reconstitution of SA biosynthesis in *N. benthamiana* and the untargeted plant metabolomics. V.C., J.-K.W. performed the genetics experiments, metabolomic profiling, and the pathogen inoculation experiment. A.B. and A.S. assisted with various genetic, biochemical and metabolomics experiments. C.M.G. performed chemical synthesis. T.P. performed ligand docking. M.P.T.-S. and J.-K.W. interpreted the results and wrote the paper.

## Supplemental Information for

**Supplemental Figure 1.**
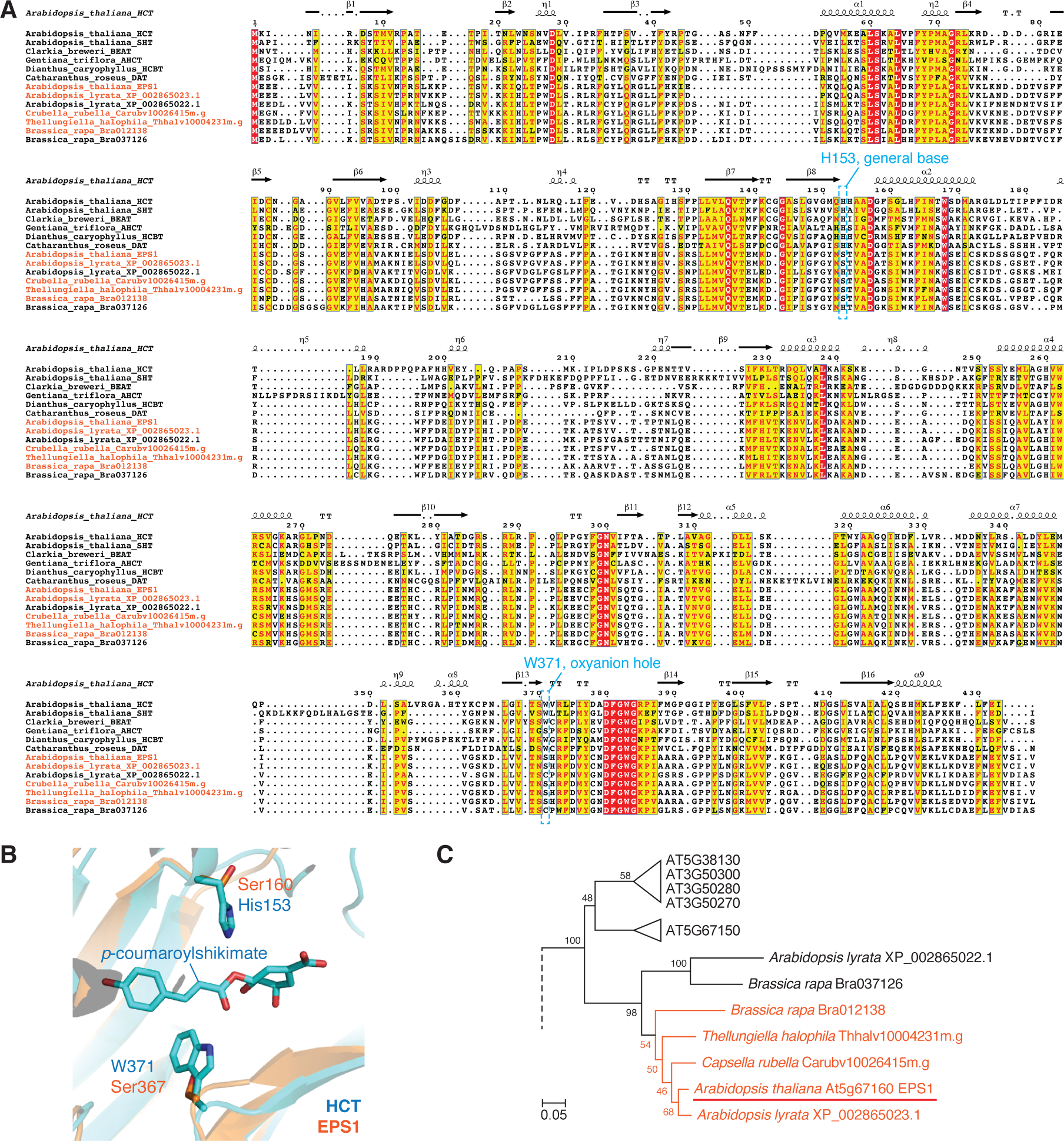
EPS1 represents a unique clade of BAHD acyltransferase family proteins associated with the Brassicaceae family plants. **(A)** Sequence alignment of EPS1 and its orthologs from other Brassicaceae species (orange, also illustrated in Supplemental Fig. 1C) together with several other related and well characterized BAHDs (D’Auria, 2006; Gou et al., 2009). The two catalytic residue positions important for canonical BAHD acyltransferase catalytic function are highlighted in blue dotted boxes. H153 (Arabidopsis HCT numbering) serves as a general base that deprotonates the hydroxyl or amine group of the acyl acceptor substrate, priming it for nucleophilic attack on the carbonyl carbon of the acyl CoA substrate (Levsh et al., 2016), and is ubiquitously conserved among canonical BAHDs. The indole nitrogen of Trp371 serves as an oxyanion hole that stabilizes the negative charge on the tetrahedral intermediate formed after the nucleophilic attack (Levsh et al., 2016). Trp371 is conserved among the majority of BAHDs. **(B)** Overlay of the active sites of Arabidopsis HCT (in complex with the product p-coumaroylshikimate, blue, PDB: 5KJU) and the EPS1 homology model (orange). EPS1 contains serine substitutions at the two catalytic residues His153 and Trp371 (AtHCT numbering) highly conserved among canonical BAHDs. **(C)** Neighbor-joining phylogenetic analysis of BAHDs from multiple reference genomes suggests that EPS1 belongs to a unique clade of BAHDs restricted in the Brassicaceae family (orange). The EPS1 orthologous clade is sister to three other BAHD clades also restricted in Brassicaceae. Bootstrap values (based on 1000 replicates) are indicated at the tree nodes. The scale measures evolutionary distances in substitutions per amino acid. The full tree is presented in Supplemental Data 4 and the sequence alignment is presented in Supplemental Data 5.

**Supplemental Figure 2.**
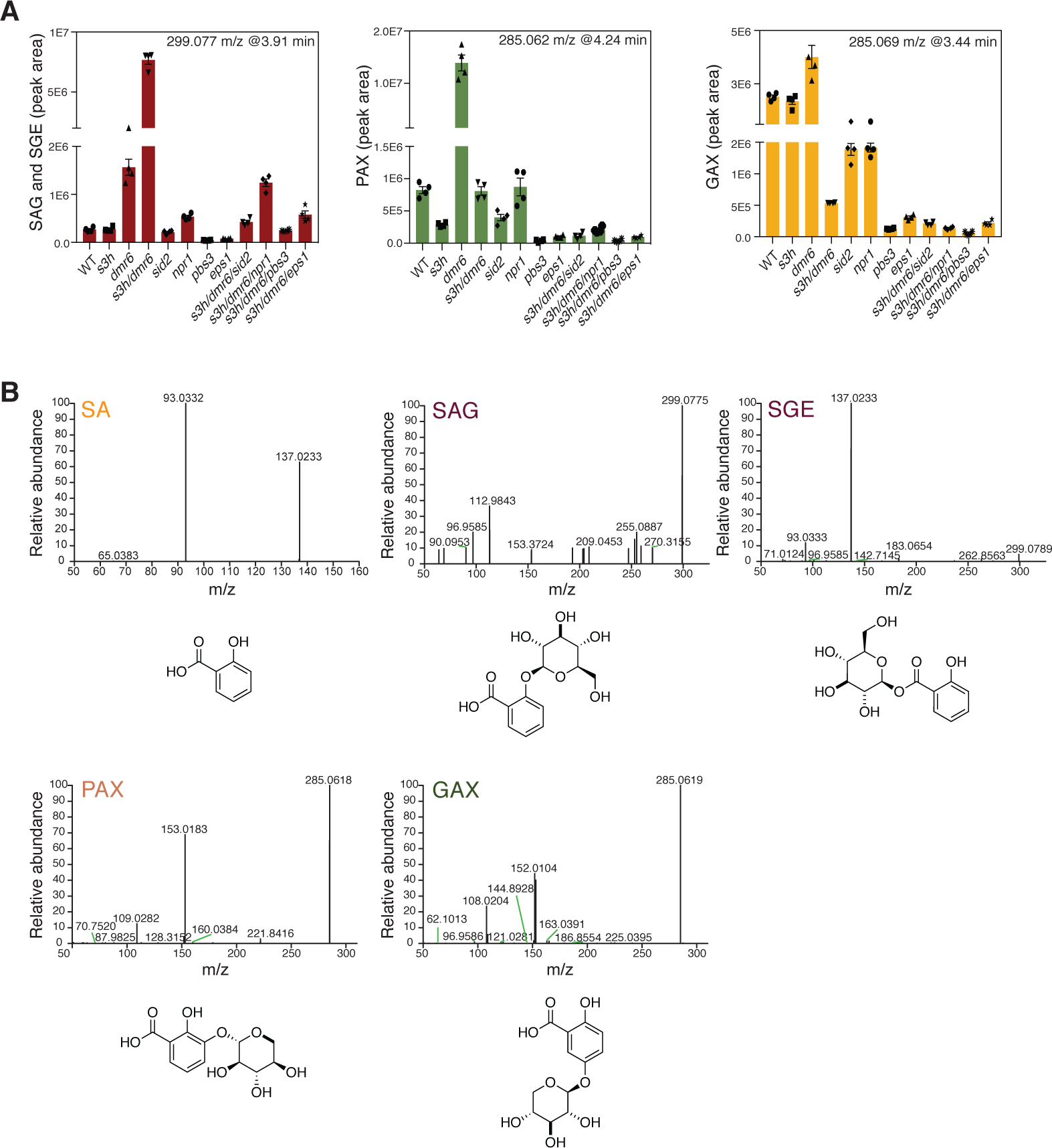
Levels of SA-derived metabolites in various Arabidopsis mutants. **(A)** Relative quantification of four major SA-derived metabolites that are differentially accumulated across various Arabidopsis SA biosynthetic and signaling mutants (n = 3). Error bars indicate standard error of the mean (SEM) based on biological triplicates. Statistical analysis was conducted by one-tailed unpaired t-test. **P < 0.01. SAG, salicylic acid 2-O-β-D-glucoside; SGE, salicylic acid glucose ester; PAX, o-pyrocatechuic acid 5-O-β-D-xyloside; GAX, gentisic acid 5-O-β-D-xyloside. The three metabolites were chromatographically separated on a C18 column, measured by LC-HRAM-MS and data processed by MZmine2. **(B)** The measured MS^2^ spectra of SA, SAG, SGE, PAX and GAX and their chemical structures.

**Supplemental Figure 3.**
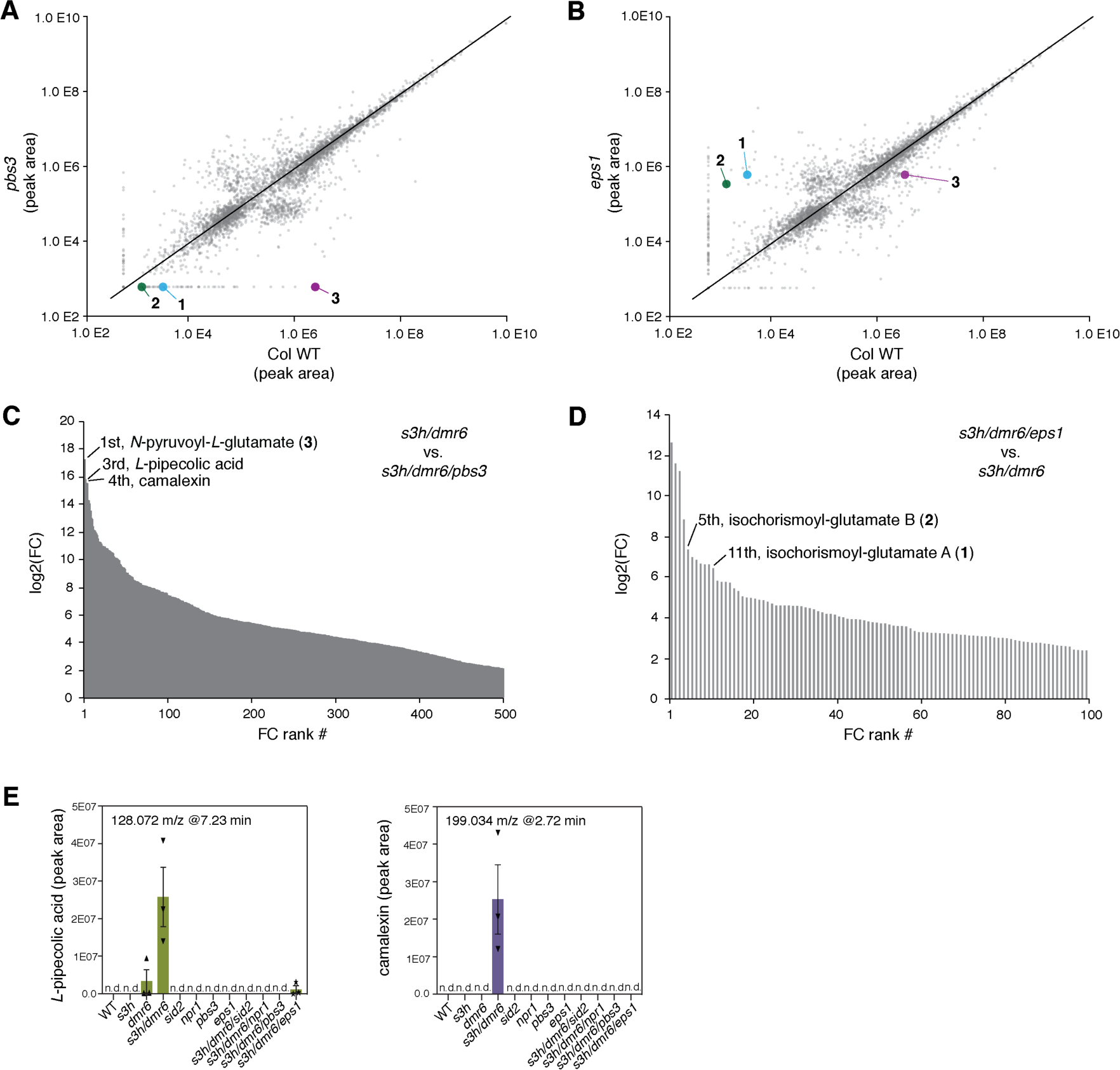
Analysis of differentially accumulated metabolic features across Arabidopsis SA-biosynthetic mutants. **(A)** Pairwise scatter plot comparing the abundance of global metabolic features of wild type and pbs3. The plot shows the accumulation of **1**-**3** in wild type, which are absent in pbs3. Data were collected using LC-HRAM-MS under negative mode. Each dot represents a metabolic feature defined by unique m/z and retention time values (n =3). **(B)** Pairwise scatter plot comparing the abundance of global metabolic features of wild type and eps1, which shows hyper-accumulation of **1** and **2** in eps1 compared to wild type (n =3). **(C)** Ranking of metabolic features that are differentially accumulated between s3h/dmr6 and s3h/dmr6/pbs3 based on Log2 fold change. **(D)** Ranking of metabolic features that are differentially accumulated between s3h/dmr6/eps1 and s3h/dmr6 based on Log2 fold change. **(E)** Relative quantification of L-pipecolic acid and camelexin that are differentially accumulated across various Arabidopsis SA biosynthetic and signaling mutants (n = 3). Error bars indicate standard error of the mean (SEM) based on biological triplicates.

**Supplemental Figure 4.**
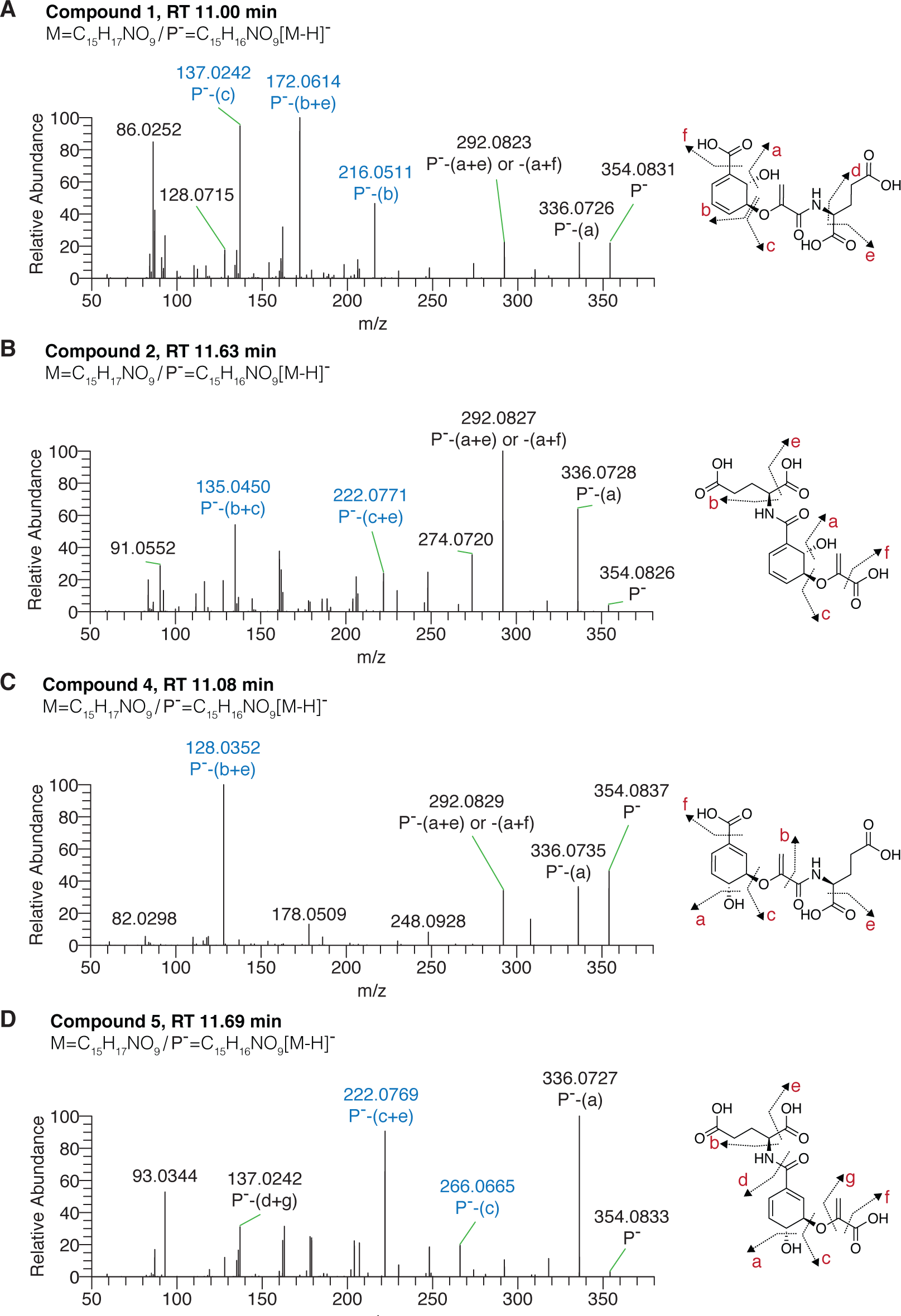
Structural elucidation of compounds **1, 2, 4** and **5**. MS^2^ fragmentation patterns of **1, 2, 4** and **5 (A-D)**. The proposed fragmentation sites are shown. P-, protonated precursor ion. The unique fragments associated with each compound that helped to determine the identity of the specific structural isomer are highlighted in blue.

**Supplemental Figure 5.**
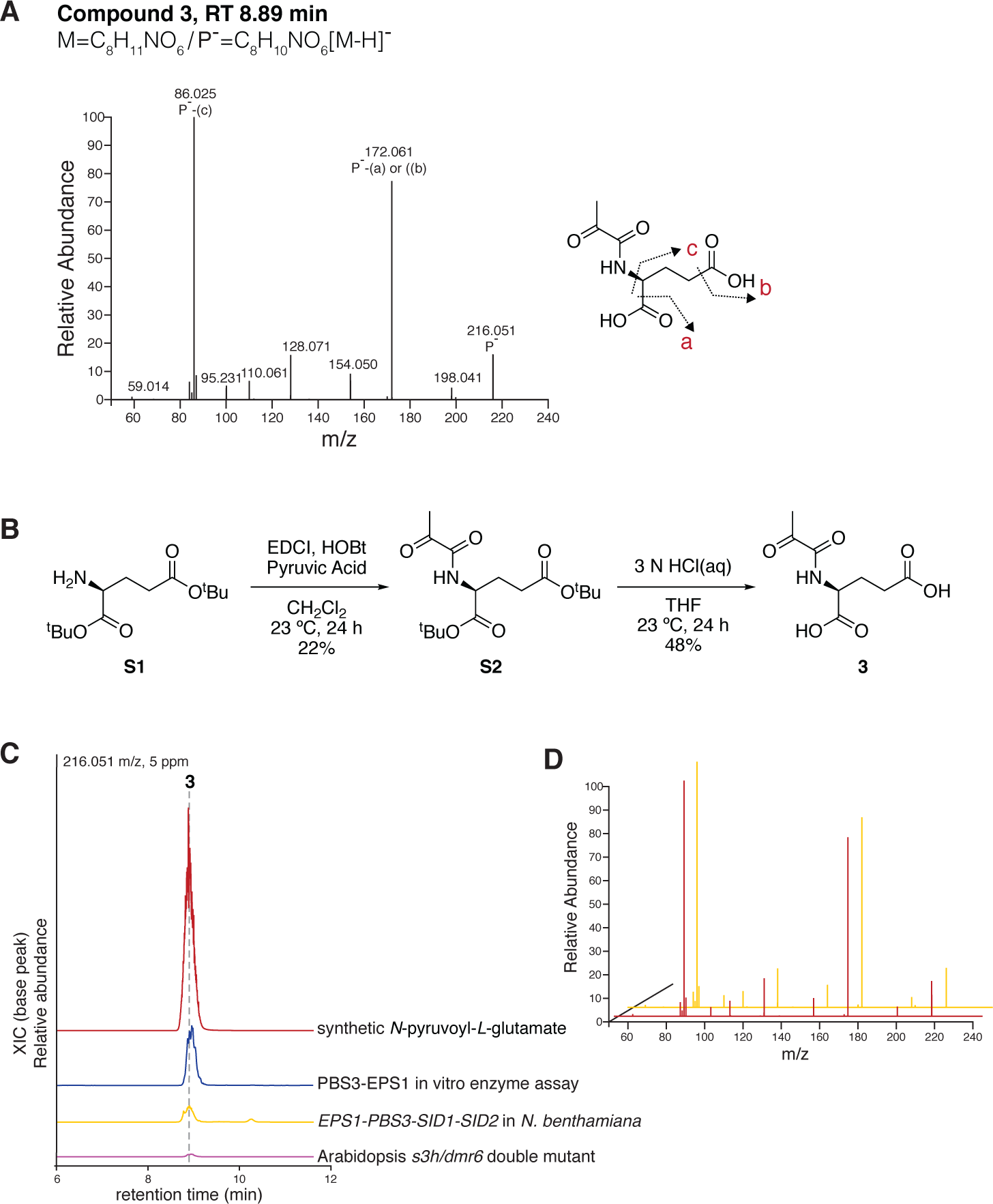
Structural elucidation of compound 3. **(A)** MS^2^ fragmentation patterns of **3**. The proposed fragmentation sites are shown. P-, protonated precursor ion. **(B)** Organic synthesis scheme for **3**. **(C)** Overlay of extracted ion chromatogram (XICs) showing the detection of **3** from LC-HRAM-MS analysis of synthetic **3**, representative PBS3-EPS1 enzyme assay, *N. benthamiana* co-expressing *EPS1-PBS3-SID1-SID2* and the Arabidopsis *s3h/dmr6* double mutant. **(D)** The MS^2^ spectrum match between synthetic **3** and **3** from *N. benthamiana* co-expressing *EPS1-PBS3-SID1-SID2* is shown.

**Supplemental Figure 6.**
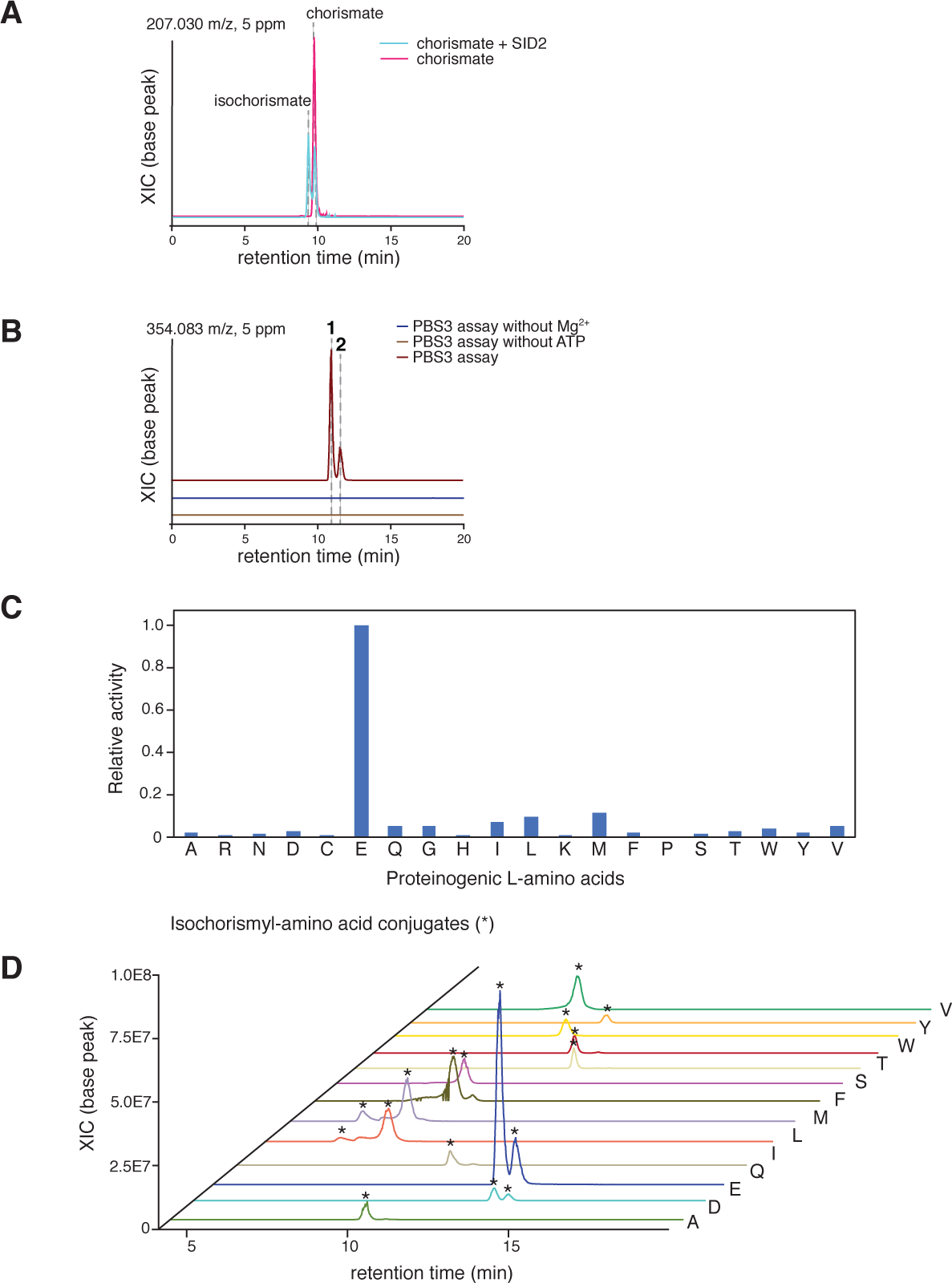
Characterization of the biochemical activity of PBS3 *in vitro*. **(A)** A preassay using SID2 to enzymatically generate isochorismate. 2 µg of SID2 was added a 100 µL reaction containing 50 mM Tris, 1 mM *L-*glutamate, 5 mM chorismate, 5 mM ATP, 5 mM MgCl_2_. Incubation for 5 minutes at 25 degrees resulted in the production of isochorismate. The reaction also reaches equilibrium that gives approximately equal molar of isochorismate and chorismate. **(B)** PBS3 activity requires ATP and Mg^2+^. **(C)** Amino acid preference of PBS3. Individual assays were carried out using isochorismate as the acyl substrate and 1 mM of each of the 20 proteinogenic amino acids as the amino acid substrate. The relative activity was determined based on AMP production. **(D)** The production of isochorismoyl-amino acid putatively identified as [M-H]^-^ ions by LC-HRAM-MS analysis are denoted by asterisks. The production of various isochorismoyl-amino acid conjugates was also queried by analyzing the MS data for [M-H]^-^ ions of the potential isochrismyol conjugate with the 20 proteinogenic *L*-amino acids: *L*-alanine (296.077 m/z, 5 ppm), *L*-aspartic acid (340.067 m/z, 5ppm), *L*-glutamic acid (354.083 m/z, 5ppm), *L*-glutamine (353.099 m/z, 5ppm), *L*-isoleucine (338.125 m/z, 5ppm), *L*-leucine (338.125 m/z, 5ppm), *L*-methionine (356.081 m/z, 5ppm), *L*-phenylalanine (372.108 m/z, 5ppm), *L*-serine (312.072 m/z, 5ppm), *L*-threonine (326.087 m/z, 5ppm), *L*-tryptophan (411.119 m/z, 5ppm), *L*-tyrosine (388.104 m/z, 5ppm), and *L*-valine (324.109 m/z, 5ppm).

**Supplemental Figure 7.**
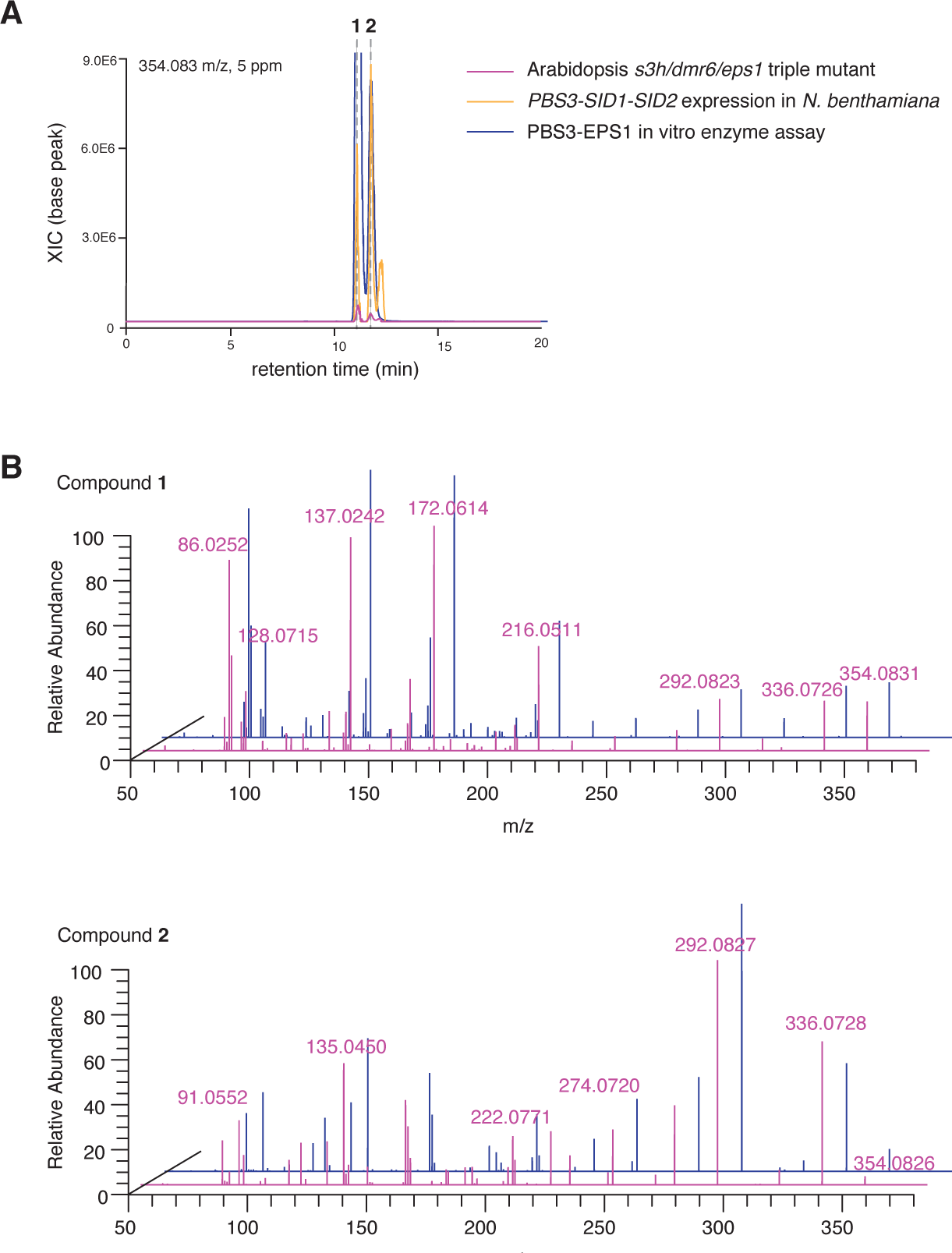
Compound 1 and 2 identity verification across Arabidopsis, N. benthamiana and in vitro enzyme assay experiments. **(A)** Overlay of extracted ion chromatogram (XICs) showing the detection of **1** and **2** from LC-HRAM-MS analyses of representative PBS3-EPS1 enzyme assay, N. benthamiana co-expressing PBS3-SID1-SID2 and the Arabidopsis s3h/dmr6/eps1 triple mutant. **(B)** The MS^2^ spectra of **1** and **2** measured from the Arabidopsis s3h/dmr6/eps1 triple mutant sample and the PBS3-EPS1 enzyme assay, which are overlaid.

**Supplemental Figure 8.**
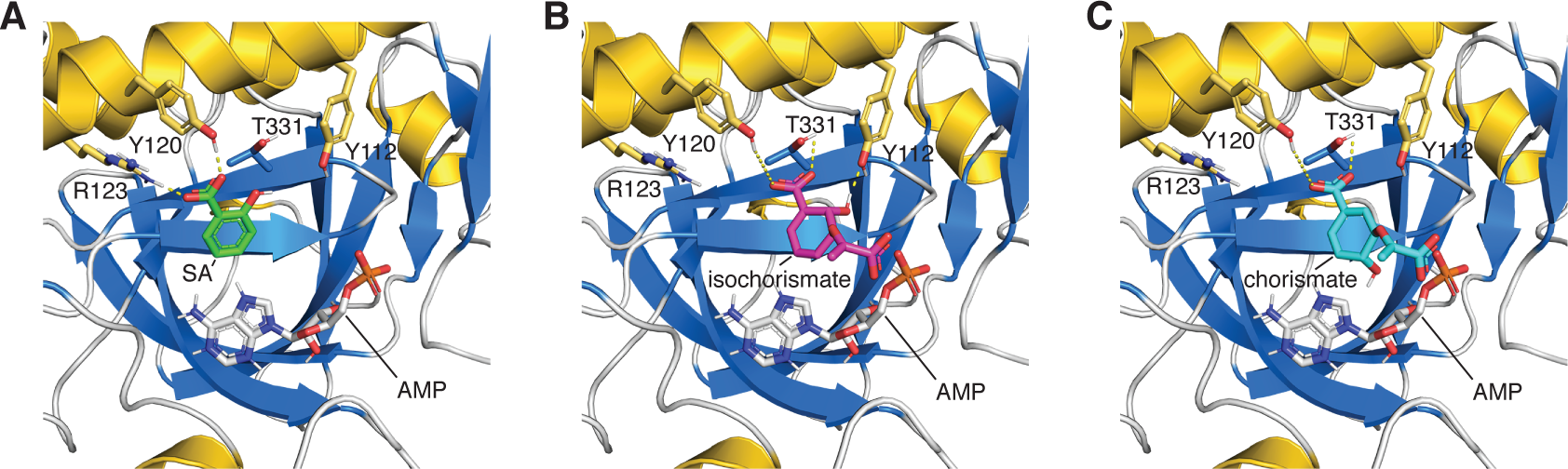
Simulated docking of isochorismate and chorismate to the PBS3 crystal structure. **(A)** The crystal structure of PBS3 in complex with SA and AMP (PDB: 4EQL) showing the catalytically nonproductive binding mode of SA in the active site. **(B)** The top simulated binding mode of isochorismate in the active site with the calculated binding affinity of −6.9 kcal/mol. **(C)** The top simulated binding mode of chorismate in the active site with the calculated binding affinity of −6.8 kcal/mol. The 0.1 kcal/mol decrease in binding affinity compared to isochorismate is likely due to the differential placement of the 4-hydroxyl in chorismate. In contrast, the 2-hydroxyl engages hydrogen bonding with the p-hydroxyl of Y112 **(B)**. The 4-hydroxyl of chorismate may also impede ATP binding. Active-site residues potentially involved in substrate binding are labeled.

**Supplemental Figure 9.**
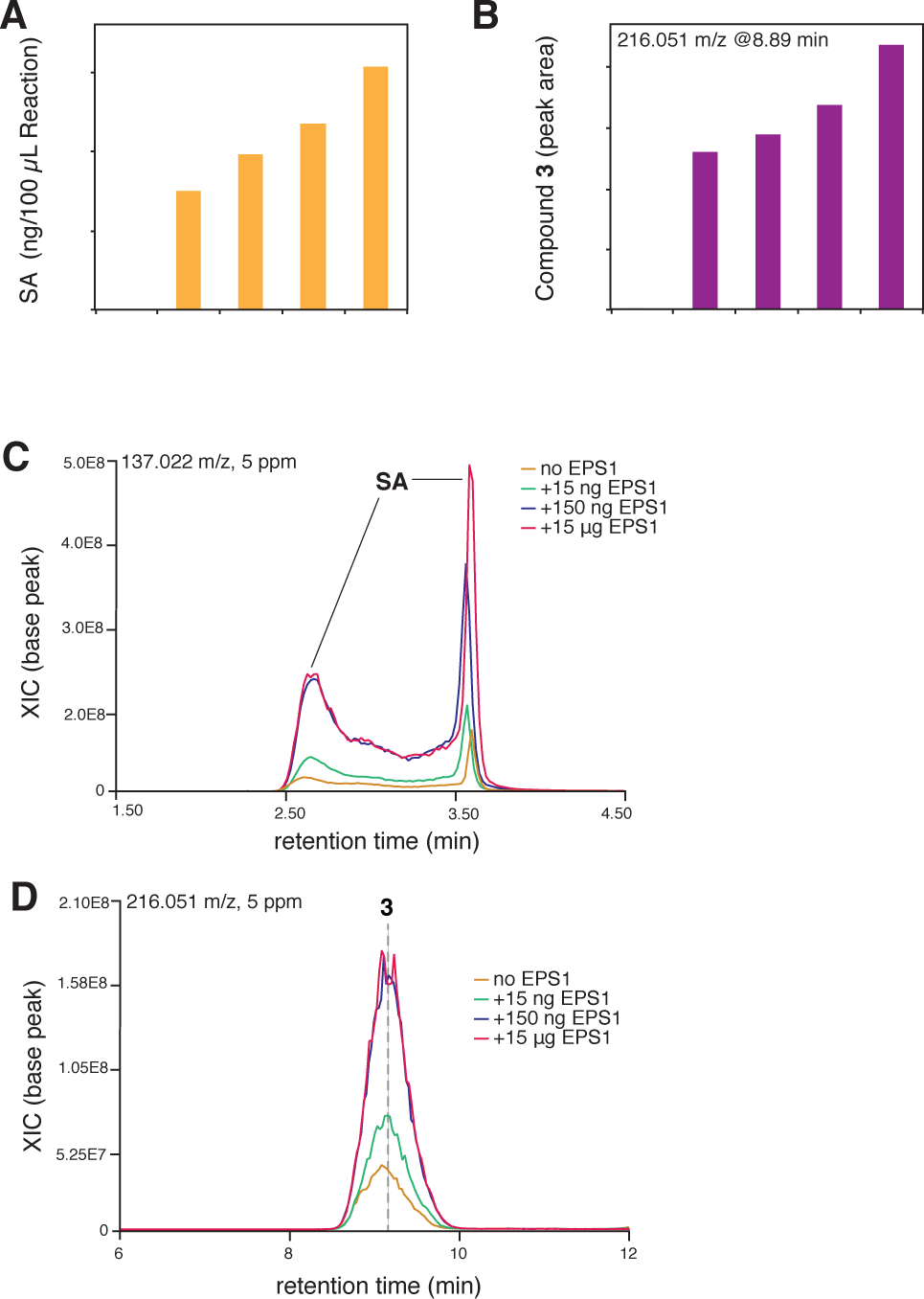
EPS1 enhances the rate of SA production from **1**. **(A)** SA production in PBS3 enzyme assays from spontaneous decay of **1**, with an estimated rate of 0.21 pkat. **(B)** Production of 3 in PBS3 enzyme assays from spontaneous decay of **1**. **(C)** Representative extracted ion chromatograms (XICs) showing EPS1-dependent production of SA from **1**. The chromatograms shown were were resolved using normal-phase chromatography. SA splits into two peaks under such chromatographic condition. **(D)** Representative XICs showing EPS1-dependent production of **3** from **1**. Also see Fig. 3D for the EPS1-dependent depletion of **1**-**2**.

**Supplemental Figure 10.**
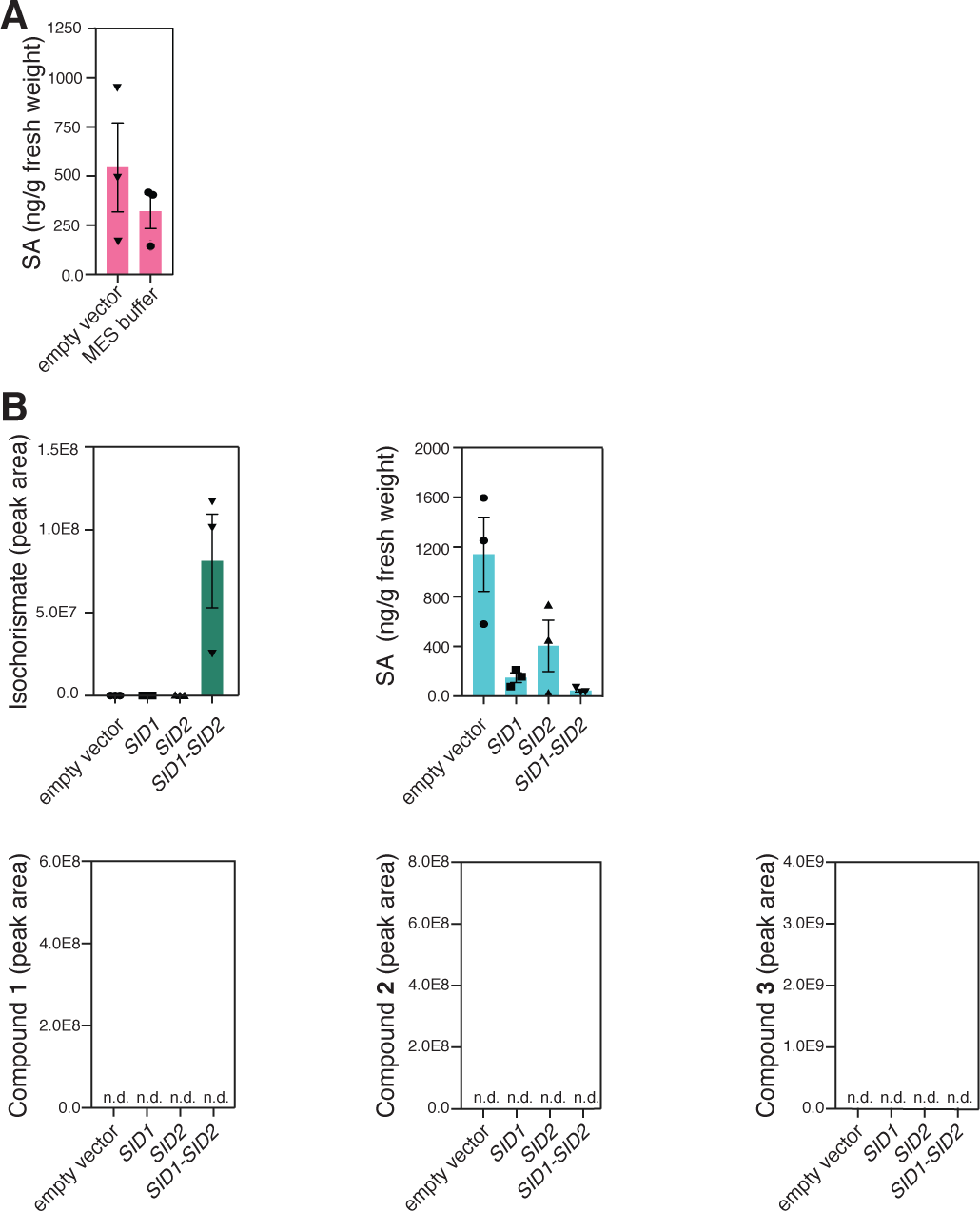
Supplementary metabolite quantification data from the N. benthamiana infiltration experiments. **(A)** Infiltration of Agrobacterium carrying empty vector does not induce de novo SA biosynthesis compared to the MES buffer control. **(B)** Quantification of isochorismate, SA and compound **1**-**3** in *SID1, SID2* and *SID1-SID2* infiltration experiments in N. benthamiana. Error bars indicate SEM based on biological triplicates (n = 3). n.d., not detected.

**Supplemental Figure 11.**
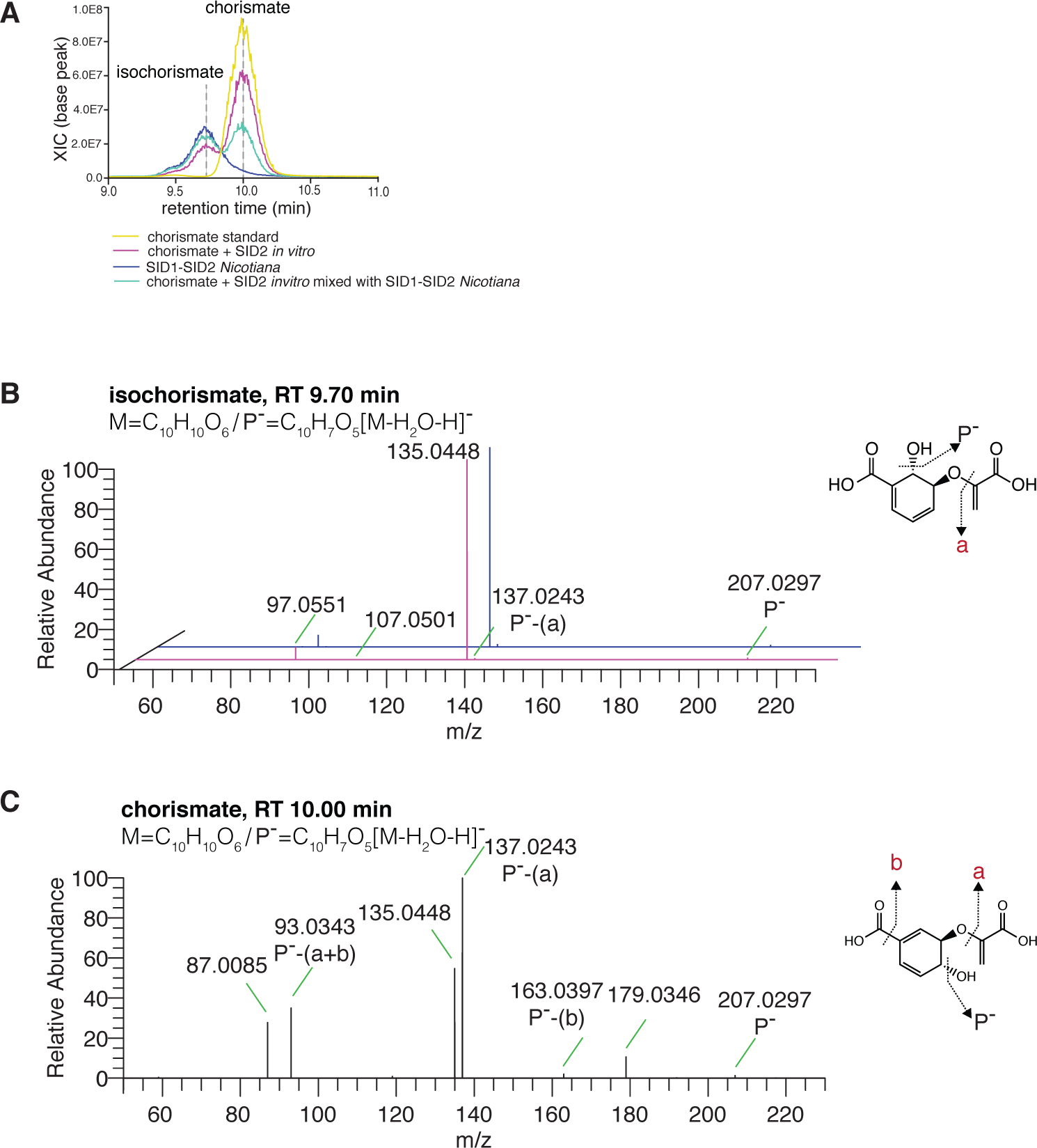
LC-HRAM-MS-based detection of isochorismate in plant extracts. **(A)** Representative extracted ion chromatograms (XICs) showing the peaks corresponding to isochorismate and chorismate from various samples. **(B)** The MS^2^ spectra of isochorismate measured from the plant extract and the SID2 enzyme assay, which are overlaid. **(C)** The MS^2^ spectrum of chorismate standard. P-, protonated precursor ion.

**Supplemental Figure 12.**
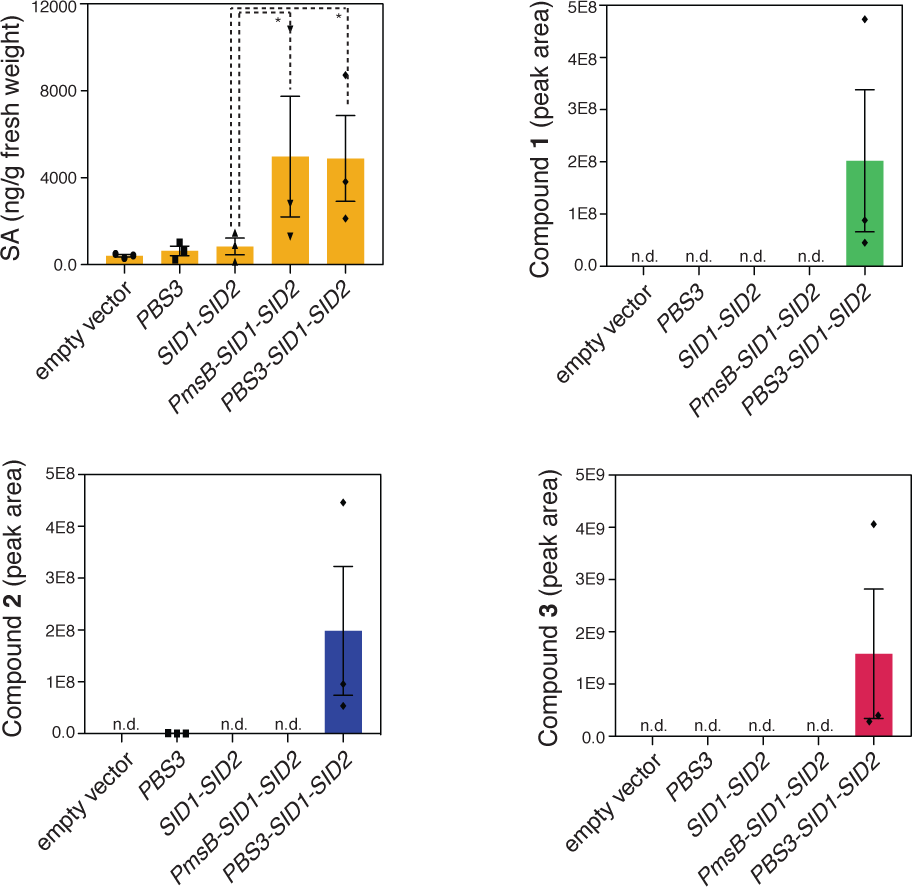
Reconstituted SA biosynthesis in N. benthamiana by Agrobacterium-mediated transient coexpression. Due to the variable nature of reconstituting SA biosynthesis in N. benthamiana, we show data that were derived from an independent experiment from data shown in Fig. 4. Quantification SA and compounds **1**-**3** in infiltrated N. benthamiana (n = 3) was shown. Error bars indicate SEM based on biological triplicates. n.d., not detected. Statistical analysis was conducted by one-tailed Mann-Whitney test. *P < 0.05.

## References

1. Alonso, J. M. (2003). Genome-Wide Insertional Mutagenesis of Arabidopsis thaliana. Science 301:653–657.

2. Amick Dempsey, D. (1999). Salicylic Acid and Disease Resistance in Plants. CRC Crit. Rev. Plant Sci. 18:547–575.

3. Baird, D. C. (1995). Experimentation: An Introduction to Measurement Theory and Experiment Design. Prentice-Hall.

4. Cao, H., Glazebrook, J., Clarke, J. D., Volko, S., and Dong, X. (1997). The Arabidopsis NPR1 gene that controls systemic acquired resistance encodes a novel protein containing ankyrin repeats. Cell 88:57–63.

5. Chen, Z., Zheng, Z., Huang, J., Lai, Z., and Fan, B. (2009). Biosynthesis of salicylic acid in plants. Plant Signal. Behav. 4:493–496.

6. Chong, J., Soufan, O., Li, C., Caraus, I., Li, S., Bourque, G., Wishart, D. S., and Xia, J. (2018). MetaboAnalyst 4.0: towards more transparent and integrative metabolomics analysis. Nucleic Acids Res. 46:W486–W494.

7. Dempsey, D. A., Corina Vlot, A., Wildermuth, M. C., and Klessig, D. F. (2011). Salicylic Acid Biosynthesis and Metabolism. The Arabidopsis Book 9:e0156.

8. Ding, Y., Sun, T., Ao, K., Peng, Y., Zhang, Y., Li, X., and Zhang, Y. (2018). Opposite Roles of Salicylic Acid Receptors NPR1 and NPR3/NPR4 in Transcriptional Regulation of Plant Immunity. Cell 173:1454–1467.e15.

9. Di Tommaso, P., Moretti, S., Xenarios, I., Orobitg, M., Montanyola, A., Chang, J.-M., Taly, J.-F., and Notredame, C. (2011). T-Coffee: a web server for the multiple sequence alignment of protein and RNA sequences using structural information and homology extension. Nucleic Acids Res. 39:W13–7.

10. Edgar, R. C. (2004). MUSCLE: multiple sequence alignment with high accuracy and high throughput. Nucleic Acids Research 32:1792–1797.

11. Hoekema, A., Hirsch, P. R., Hooykaas, P. J. J., and Schilperoort, R. A. (1983). A binary plant vector strategy based on separation of vir- and T-region of the Agrobacterium tumefaciens Ti-plasmid. Nature 303:179.

12. Huang, J., Gu, M., Lai, Z., Fan, B., Shi, K., Zhou, Y.-H., Yu, J.-Q., and Chen, Z. (2010). Functional analysis of the Arabidopsis PAL gene family in plant growth, development, and response to environmental stress. Plant Physiol. 153:1526–1538.

13. Jagadeeswaran, G., Raina, S., Acharya, B. R., Maqbool, S. B., Mosher, S. L., Appel, H. M., Schultz, J. C., Klessig, D. F., and Raina, R. (2007). Arabidopsis GH3-LIKE DEFENSE GENE 1 is required for accumulation of salicylic acid, activation of defense responses and resistance to Pseudomonas syringae. Plant J. 51:234–246.

14. Katagiri, F., Thilmony, R., and He, S. Y. (2002). The Arabidopsis Thaliana-Pseudomonas Syringae Interaction. Arabidopsis Book 1:e0039.

15. Kleinboelting, N., Huep, G., Kloetgen, A., Viehoever, P., and Weisshaar, B. (2012). GABI-Kat SimpleSearch: new features of the Arabidopsis thaliana T-DNA mutant database. Nucleic Acids Res. 40:D1211–5.

16. Koji, O., Yoichiro, S., Akira, C., Sayuri, F., Misato, N., and Masataka, S. (2008). Amino acid derivative. Patent Advance Access published July 10, 2008.

17. Kumar, S., Stecher, G., and Tamura, K. (2016). MEGA7: Molecular Evolutionary Genetics Analysis Version 7.0 for Bigger Datasets. Mol. Biol. Evol. 33:1870–1874.

18. Lee, M. W., Lu, H., Jung, H. W., and Greenberg, J. T. (2007). A key role for the Arabidopsis WIN3 protein in disease resistance triggered by Pseudomonas syringae that secrete AvrRpt2. Mol. Plant. Microbe. Interact. 20:1192–1200.

19. León, J., Shulaev, V., Yalpani, N., Lawton, M. A., and Raskin, I. (1995). Benzoic acid 2-hydroxylase, a soluble oxygenase from tobacco, catalyzes salicylic acid biosynthesis. Proc. Natl. Acad. Sci. U. S. A. 92:10413–10417.

20. Levsh, O., Chiang, Y.-C., Tung, C. F., Noel, J. P., Wang, Y., and Weng, J.-K. (2016). Dynamic Conformational States Dictate Selectivity toward the Native Substrate in a Substrate-Permissive Acyltransferase. Biochemistry 55:6314–6326.

21. Nawrath, C., Heck, S., Parinthawong, N., and Métraux, J.-P. (2002). EDS5, an essential component of salicylic acid-dependent signaling for disease resistance in Arabidopsis, is a member of the MATE transporter family. Plant Cell 14:275–286.

22. Nobuta, K., Okrent, R. A., Stoutemyer, M., Rodibaugh, N., Kempema, L., Wildermuth, M. C., and Innes, R. W. (2007). The GH3 acyl adenylase family member PBS3 regulates salicylic acid-dependent defense responses in Arabidopsis. Plant Physiol. 144:1144–1156.

23. Okrent, R. A., and Wildermuth, M. C. (2011). Evolutionary history of the GH3 family of acyl adenylases in rosids. Plant Mol. Biol. 76:489–505.

24. Okrent, R. A., Brooks, M. D., and Wildermuth, M. C. (2009). Arabidopsis GH3.12 (PBS3) conjugates amino acids to 4-substituted benzoates and is inhibited by salicylate. J. Biol. Chem. 284:9742–9754.

25. Pallas, J. A., Paiva, N. L., Lamb, C., and Dixon, R. A. (1996). Tobacco plants epigenetically suppressed in phenylalanine ammonia-lyase expression do not develop systemic acquired resistance in response to infection by tobacco mosaic virus. The Plant Journal 10:281–293.

26. Park, J.-E., Park, J.-Y., Kim, Y.-S., Staswick, P. E., Jeon, J., Yun, J., Kim, S.-Y., Kim, J., Lee, Y.-H., and Park, C.-M. (2007). GH3-mediated auxin homeostasis links growth regulation with stress adaptation response in Arabidopsis. J. Biol. Chem. 282:10036–10046.

27. Pluskal, T., Castillo, S., Villar-Briones, A., and Orešič, M. (2010). MZmine 2: Modular framework for processing, visualizing, and analyzing mass spectrometry-based molecular profile data. BMC Bioinformatics 11.

28. Raskin, I. (1992). Role of Salicylic Acid in Plants. Annu. Rev. Plant Physiol. Plant Mol. Biol. 43:439–463.

29. Rekhter, D., Lüdke, D., Ding, Y., Feussner, K., Zienkiewicz, K., Lipka, V., Wiermer, M., Zhang, Y., and Feussner, I. (2019). Isochorismate-derived biosynthesis of the plant stress hormone salicylic acid. Science 365:498–502.

30. Robert, X., and Gouet, P. (2014). Deciphering key features in protein structures with the new ENDscript server. Nucleic Acids Res. 42:W320–4.

31. Sainsbury, F., Thuenemann, E. C., and Lomonossoff, G. P. (2009). pEAQ: versatile expression vectors for easy and quick transient expression of heterologous proteins in plants. Plant Biotechnol. J. 7:682–693.

32. Trott, O., and Olson, A. J. (2010). AutoDock Vina: improving the speed and accuracy of docking with a new scoring function, efficient optimization, and multithreading. J. Comput. Chem. 31:455–461.

33. van Damme, M., Huibers, R. P., Elberse, J., and Van den Ackerveken, G. (2008). Arabidopsis DMR6 encodes a putative 2OG-Fe(II) oxygenase that is defense-associated but required for susceptibility to downy mildew. Plant J. 54:785–793.

34. Verberne, M. C., Verpoorte, R., Bol, J. F., Mercado-Blanco, J., and Linthorst, H. J. (2000). Overproduction of salicylic acid in plants by bacterial transgenes enhances pathogen resistance. Nat. Biotechnol. 18:779–783.

35. Waterhouse, A., Bertoni, M., Bienert, S., Studer, G., Tauriello, G., Gumienny, R., Heer, F. T., de Beer, T. A. P., Rempfer, C., Bordoli, L., et al. (2018). SWISS-MODEL: homology modelling of protein structures and complexes. Nucleic Acids Res. 46:W296–W303.

36. Wehbe, J., Rolland, V., Fruchier, A., Roumestant, M.-L., and Martinez, J. (2004). Enantioselective synthesis of new 4-substituted glutamic acid derivatives. Tetrahedron: Asymmetry 15:851–858.

37. Weng, J.-K., and Noel, J. P. (2012). The remarkable pliability and promiscuity of specialized metabolism. Cold Spring Harb. Symp. Quant. Biol. 77:309–320.

38. Westfall, C. S., Zubieta, C., Herrmann, J., Kapp, U., Nanao, M. H., and Jez, J. M. (2012). Structural basis for prereceptor modulation of plant hormones by GH3 proteins. Science 336:1708–1711.

39. Westfall, C. S., Sherp, A. M., Zubieta, C., Alvarez, S., Schraft, E., Marcellin, R., Ramirez, L., and Jez, J. M. (2016). Arabidopsis thaliana GH3.5 acyl acid amido synthetase mediates metabolic crosstalk in auxin and salicylic acid homeostasis. Proc. Natl. Acad. Sci. U. S. A. 113:13917–13922.

40. Wildermuth, M. C., Dewdney, J., Wu, G., and Ausubel, F. M. (2001). Isochorismate synthase is required to synthesize salicylic acid for plant defence. Nature 414:562–565.

41. Wu, Y., Zhang, D., Chu, J. Y., Boyle, P., Wang, Y., Brindle, I. D., De Luca, V., and Després, C. (2012). The Arabidopsis NPR1 Protein Is a Receptor for the Plant Defense Hormone Salicylic Acid. Cell Rep. 1:639–647.

42. Zeilmaker, T., Ludwig, N. R., Elberse, J., Seidl, M. F., Berke, L., Van Doorn, A., Schuurink, R. C., Snel, B., and Van den Ackerveken, G. (2015). DOWNY MILDEW RESISTANT 6 and DMR6-LIKE OXYGENASE 1 are partially redundant but distinct suppressors of immunity in Arabidopsis. Plant J. 81:210–222.

43. Zhang, K., Halitschke, R., Yin, C., Liu, C.-J., and Gan, S.-S. (2013). Salicylic acid 3-hydroxylase regulates Arabidopsis leaf longevity by mediating salicylic acid catabolism. Proc. Natl. Acad. Sci. U. S. A. 110:14807–14812.

44. Zhang, Y., Zhao, L., Zhao, J., Li, Y., Wang, J., Guo, R., Gan, S., Liu, C.-J., and Zhang, K. (2017). S5H/DMR6 Encodes a Salicylic Acid 5-Hydroxylase That Fine-Tunes Salicylic Acid Homeostasis. Plant Physiol. 175:1082–1093.

45. Zheng, Z., Qualley, A., Fan, B., Dudareva, N., and Chen, Z. (2009). An important role of a BAHD acyl transferase-like protein in plant innate immunity. Plant J. 57:1040–1053.

46. Zheng, Z., Guo, Y., Novák, O., Chen, W., Ljung, K., Noel, J. P., and Chory, J. (2016). Local auxin metabolism regulates environment-induced hypocotyl elongation. Nat Plants 2:16025.

47. Zhou, Y., Memelink, J., and Linthorst, H. J. M. (2018). An E. coli biosensor for screening of cDNA libraries for isochorismate pyruvate lyase-encoding cDNAs. Mol. Genet. Genomics 293:1181–1190.

## Supplemental references

1. D’Auria, J. C. (2006). Acyltransferases in plants: a good time to be BAHD. Curr. Opin. Plant Biol. 9:331–340.

2. Gou, J.-Y., Yu, X.-H., and Liu, C.-J. (2009). A hydroxycinnamoyltransferase responsible for synthesizing suberin aromatics in Arabidopsis. Proc. Natl. Acad. Sci. U. S. A. 106:18855–18860.

3. Levsh, O., Chiang, Y.-C., Tung, C. F., Noel, J. P., Wang, Y., and Weng, J.-K. (2016). Dynamic Conformational States Dictate Selectivity toward the Native Substrate in a Substrate-Permissive Acyltransferase. Biochemistry 55:6314–6326.

